# Complete Disruption of Autism-Susceptibility Genes by Gene-Editing Predominantly Reduces Functional Connectivity of Isogenic Human Neurons

**DOI:** 10.1101/344234

**Authors:** Eric Deneault, Sean H. White, Deivid C. Rodrigues, P. Joel Ross, Muhammad Faheem, Kirill Zaslavsky, Zhuozhi Wang, Roumiana Alexandrova, Giovanna Pellecchia, Wei Wei, Alina Piekna, Gaganjot Kaur, Jennifer L. Howe, Vickie Kwan, Bhooma Thiruvahindrapuram, Susan Walker, Peter Pasceri, Daniele Merico, Ryan K.C. Yuen, Karun K. Singh, James Ellis, Stephen W. Scherer

## Abstract

Autism Spectrum Disorder is phenotypically and genetically heterogeneous, but genomic analyses have identified candidate susceptibility genes. We present a CRISPR gene editing strategy to insert a protein tag and premature termination sites creating an induced pluripotent stem cell (iPSC) knockout resource for functional studies of 10 ASD-relevant genes (*AFF2/FMR2, ANOS1, ASTN2, ATRX*, *CACNA1C*, *CHD8, DLGAP2, KCNQ2*, *SCN2A*, *TENM1*). Neurogenin 2 (NEUROG2)-directed differentiation of iPSCs allowed production of cortical excitatory neurons, and mutant proteins were not detectable. RNAseq revealed convergence of several neuronal networks. Using both patch-clamp and multi-electrode array approaches, the electrophysiological deficits measured were distinct for different mutations. However, they culminated in a consistent reduction in synaptic activity, including reduced spontaneous excitatory post-synaptic current frequencies in *AFF2/FMR2-*, *ASTN2-, ATRX*-, *KCNQ2*- and *SCN2A*-null neurons. Despite ASD susceptibility genes belonging to different gene ontologies, isogenic stem cell resources can reveal common functional phenotypes, such as reduced functional connectivity.

## Introduction

Autism spectrum disorder (ASD) is a lifelong neurodevelopmental condition affecting reciprocal social interaction and communication, accompanied with restricted and repetitive behaviours (DSM-V, 2013). Familial clustering of ASD and related subclinical traits has been described, and with sibling recurrence risk estimates ranging from 8.1 to 18.7 (Gronborg et al., 2013; Ozonoff et al., 2011; Risch et al., 2014), a significant amount of familial liability is attributed to genetic factors (Colvert et al., 2015). Genomic microarray and sequencing studies have identified that ∼10% of individuals have an identifiable genetic condition, and in fact, there are over 100 genetic disorders that can exhibit features of ASD, e.g., Fragile X and Rett syndromes (Betancur, 2011). Dozens of additional penetrant susceptibility genes have also been implicated in ASD (De Rubeis et al., 2014; Gilman et al., 2011; Pinto et al., 2014; Tammimies et al., 2015; Yuen et al., 2017), some being used in clinical diagnostic testing settings (Carter and Scherer, 2013). Genetically-identified ASD risk genes are enriched in broader functional groups consisting of synapse function, RNA processing, and transcriptional regulation (Bourgeron, 2015; De Rubeis et al., 2014; Geschwind and State, 2015; Pinto et al., 2014; Yuen et al., 2017; Yuen et al., 2016). Importantly, so far, each risk gene or copy number variation (CNV) implicated in ASD accounts for <1% of cases, suggesting significant genetic heterogeneity (Yuen et al., 2017). Even within families, siblings can carry different penetrant mutations (Geschwind and State, 2015; Yuen et al., 2015).

Until recently, post-mortem brains were the only source of human cortical neurons to study directly the mechanistic and functional roles of ASD candidate genes *in vitro* (Varghese et al., 2017; Wintle et al., 2011). However, the still small numbers of biobanked brains, and the heterogeneous cellular content of the organ itself, as well as issues of cell viability and health are all difficult to properly control for a such complex disorder (Anagnostou et al., 2014; de la Torre-Ubieta et al., 2016). The terminal differentiation status of mature neurons also precludes any potential *in vitro* studies in particular for early-onset conditions like ASD. Alternatively, in a different approach, mature somatic cells can be reprogrammed into induced pluripotent stem cells (iPSCs) that can grow indefinitely *in vitro* (Takahashi et al., 2007; Yu et al., 2007). Such patient-specific iPSCs provide a newfound ability to study developmental processes, and functional characteristics, directly. Importantly, differentiation of human iPSCs into forebrain glutamatergic neurons may lead to model systems that recapitulate early molecular events in the trajectory of ASD development (Habela et al., 2016; Moretto et al., 2017). Directed differentiation into excitatory neurons can be achieved with high efficiency using transient ectopic expression of the transcription factor NEUROG2 (Zhang et al., 2013).

Here, we devised a precise clustered regularly interspaced short palindromic repeats (CRISPR)-based strategy to efficiently generate complete knockout (KO) of ASD-relevant genes, with all mutations made in the same “isogenic” human control iPSC line. We capitalized on the CRISPR/Cas9-mediated double-strand break (DSB) mechanism coupled with error-free single-stranded template repair (SSTR) pathways (Miyaoka et al., 2014), to introduce an all-reading-frame premature termination codon (named “StopTag”; **Figure 1A**) into a specific exon of a target gene, designed to prevent stable RNA/protein product from being made. We hypothesized that a cohort of isogenic KO lines carrying different ASD-risk null mutations would best minimize the confounding effects of genetic background and its impact on phenotype. We then explored excitatory neuron functional differences relevant to ASD for 10 different successfully-edited StopTag lines. We used transcriptional profiling and two complementary electrophysiological assays to measure various neuronal phenotypes.

**Figure 1.**
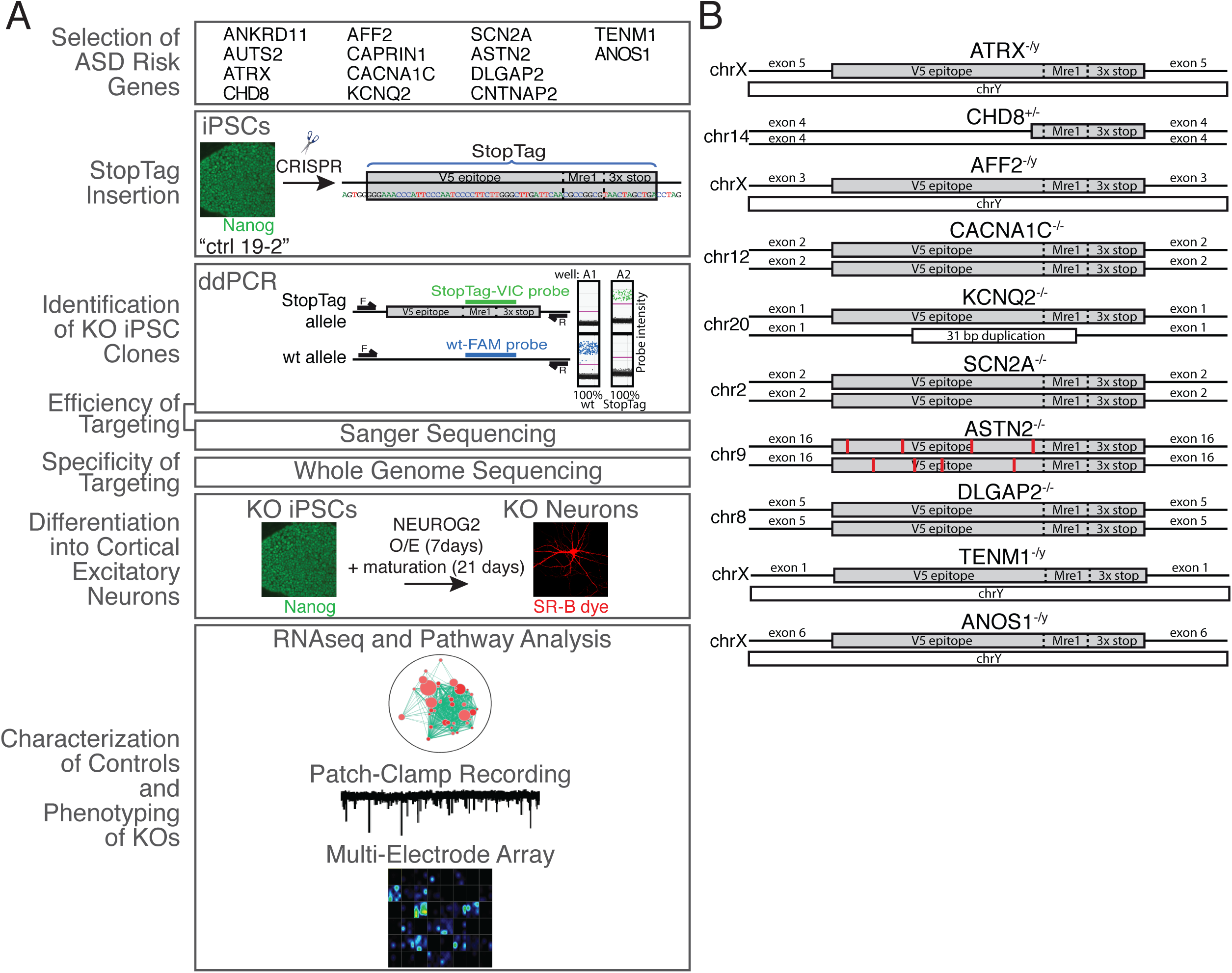
Outline of the experimental procedure to test the phenotypical consequences of gene repression in iPSC-derived glutamatergic neurons. Unaffected human iPSCs control (ctrl) 19-2, labeled in green, were subjected to CRISPR gene editing to introduce a premature termination codon, named “StopTag”, into a target exon of 14 ASD risk target genes. Knockout (KO) iPSC populations were identified by absolute quantification of StopTag versus wildtype (wt) alleles using droplet digital (ddPCR). Well A1 is an example of a cell population containing 100% wt allele (FAM signal in blue) for a given target locus, while well A2 contains 100% StopTag alleles (VIC signal in green); FAM- and VIC-associated probe sequences are presented in **Table S1**. KO iPSCs were differentiated into glutamatergic neurons, labeled in red, by means of NEUROG2 transient overexpression (O/E). Neuronal phenotypes were monitored using RNAseq, patch-clamp and multi-electrode array recordings; F and R = ddPCR primers. (B) Full-length integral StopTag sequence insertion was confirmed for all target genes except CHD8, in which the first 39 bp in 5’ of the StopTag sequence were deleted; and *ASTN2*, in which different point mutations were found (red bars). chr = chromosome; bp = base pair

Our results indicate that some ASD risk genes display reduced synaptic activity between NEUROG2-derived excitatory neurons. These data also imply that ASD genes from different classes can display the same general cellular phenotype *in vitro,* providing evidence of a common mechanism of disease and highlighting the benefits of studying ASD risk genes using an isogenic human neural system.

## Results

### Selection of ASD Risk Genes

We selected 14 candidate ASD-susceptibility genes from our ongoing whole-genome sequencing (WGS) project (the Autism Speaks MSSNG project), which aims to generate a list of penetrant genes for clinical diagnostics (Yuen et al., 2017). The evidence and priority of each gene for having a role in ASD is described in **Table 1**, as is its assignment within three different functional groupings, i.e., transcriptional regulation (*ANKRD11, AUTS2, ATRX, CHD8*), RNA processing (*AFF2/FMR2, CAPRIN1*), and synaptic and adhesion (*CACNA1C, KCNQ2, SCN2A, ASTN2, DLGAP2, CNTNAP2, TENM1, ANOS1*). The priority of a gene on such “ASD-risk gene” lists can change throughout the length of a study due to new genetic and functional data emerging (discussed below).

**Table 1.** List of 14 ASD susceptibility genes selected for this study with their corresponding functional groupings.

| Gene | Chr | Molecular Function | Module | ASD Gene Ref. | SFARI Score | OMIM Gene | Associated Disorders | ASD Syndrome |
| --- | --- | --- | --- | --- | --- | --- | --- | --- |
| *ANKRD11* | 16 | Chromatin regulation | Transcriptional regulation | (Yuen et al., 2017) | 2 | 611192 | ADHD, EPS, ID | KBG syndrome |
| *AUTS2* | 7 | Chromatin binding | Transcriptional regulation | (Yuen et al., 2017) | 3 |  | ID, DD/NDD |  |
| **ATRX** | X | Chromatin binding/helicase | Transcriptional regulation | (Brett et al., 2014) | 4 | 300032 | EP, EPS |  |
| **CHD8** | 14 | Chromatin binding/helicase | Transcriptional regulation | (Yuen et al., 2017) | 1 | 610528 | SCZ, DD/NDD, ID |  |
| **AFF2 (FMR2)** | X | RNA-binding | RNA Processing | (Yuen et al., 2017) | 4 | 300806 | ID, ASD, EPS, EP, ADHD | Fragile X syndrome |
| **CAPRIN1** | 11 | RNA-binding | RNA Processing | (Jiang et al., 2013) | 3 |  |  |  |
| **CACNA1C** | 12 | Ion channel activity | Synaptic and Adhesion | (Yuen et al., 2017) | S | 114205 | EPS, BPD, ID | Timothy syndrome |
| **KCNQ2** | 20 | Ion channel activity | Synaptic and Adhesion | (Yuen et al., 2017) | 3 | 602235 | ADHD, DD/NDD, ID |  |
| **SCN2A** | 2 | Ion channel activity | Synaptic and Adhesion | (Yuen et al., 2017) | 1 | 182390 | EPS, DD/NDD, ADHD, ID, EP |  |
| **ASTN2** | 9 | Calcium ion binding | Synaptic and Adhesion | (Lionel et al., 2014) | 3 |  | ID, EPS, DD/NDD, ADHD |  |
| **DLGAP2** | 8 | Synapse | Synaptic and Adhesion | (Marshall et al., 2008) | 4 |  |  |  |
| **CNTNAP2** | 7 | Cell adhesion | Synaptic and Adhesion | (Bakkaloglu et al., 2008) | 2S | 604569 | ADHD, EP, EPS, ID | Cortical dysplasia-focal epilepsy syndrome |
| **TENM1 (ODZ1)** | X | Cell adhesion/signal transducer | Synaptic and Adhesion | (Yuen et al., 2017) | n/a |  |  |  |
| ***ANOS1***<br>**(KAL1)** | X | Extracellular<br>matrix | Synaptic and<br>Adhesion | (Jiang et al.,<br>2013) | n/a | 300836 |  | Kallmann<br>syndrome |

Some genes like *ANOS1* and *TENM1* were considered stronger candidates for having a role in ASD early in the study but this changed as additional genetic studies were published. ADHD: attention deficit with hyperactivity disorder, EPS: extrapyramidal symptoms, ID: intellectual disability, ASD: autism spectrum disorder, EP: epilepsy, SCZ: schizophrenia, DD/NDD: developmental delay/neurodevelopmental disorder, BPD: bipolar disorder.

### StopTag Insertion into ASD Risk Genes and the Isogenic Control iPSC Line

We first used HEK293T cells to validate our SSTR-based strategy that involved introducing all-reading-frame premature termination codons (PTC; 3x stop; **Figure S1A**) into the genomic DNA corresponding to an early constitutive exon for each gene (**Table S1**), aiming for a complete expression knockout. This insertion was delivered by a synthesized single-stranded oligodeoxynucleotide (ssODN) template (**Table S1**). The inserted fragment, called “StopTag” (**
Figure 1A and S1A**), is 59 bp in length and includes a V5 epitope coding sequence and a *Mre1* restriction site. The left homology arm of ssODN was designed to insert the V5 epitope in phase with the original reading frame in order to allow the detection of truncated forms of protein, with potential residual activity, that might have escaped non-sense mediated decay (NMD) following PTC insertion. Type II CRISPR/Cas9 double-nicking (Cas9D10A) (Ran et al., 2013) system with dual guide RNA (**Table S1**) was used to reduce off-target activity. PCR amplification confirmed the integration of the StopTag within specific target loci in *CHD8*, *DLGAP2* and *KCNQ2* (**Figure S1B**).

For human iPSCs, the same StopTag insertion was used to KO each of the 14 ASD risk genes in a pluripotent and karyotypically/WGS normal iPSC line named “Ctrl 19-2” (**
Figure 1A and 2A-D**). This line was reprogrammed from an unaffected father of a child with ASD who carries a *de novo* 16p11.2 microdeletion (Marshall et al., 2008), one of the most frequent CNVs associated with autism (Kumar et al., 2008; Marshall et al., 2008; Weiss et al., 2008). Nonetheless, WGS revealed that the unaffected father did not carry any known ASD risk mutations. Enrichment of 19-2-derived KO iPSCs was based on droplet digital PCR (ddPCR) coupled with dilution culture steps (**
Figure 1A and Table S2**) (Miyaoka et al., 2014), and adapted in order to maintain the polyclonality of cellular populations during the enrichment process. On average, 4.2 enrichment plates per line were necessary to purify to 100% KO (**Table S2**).

**Figure 2.**
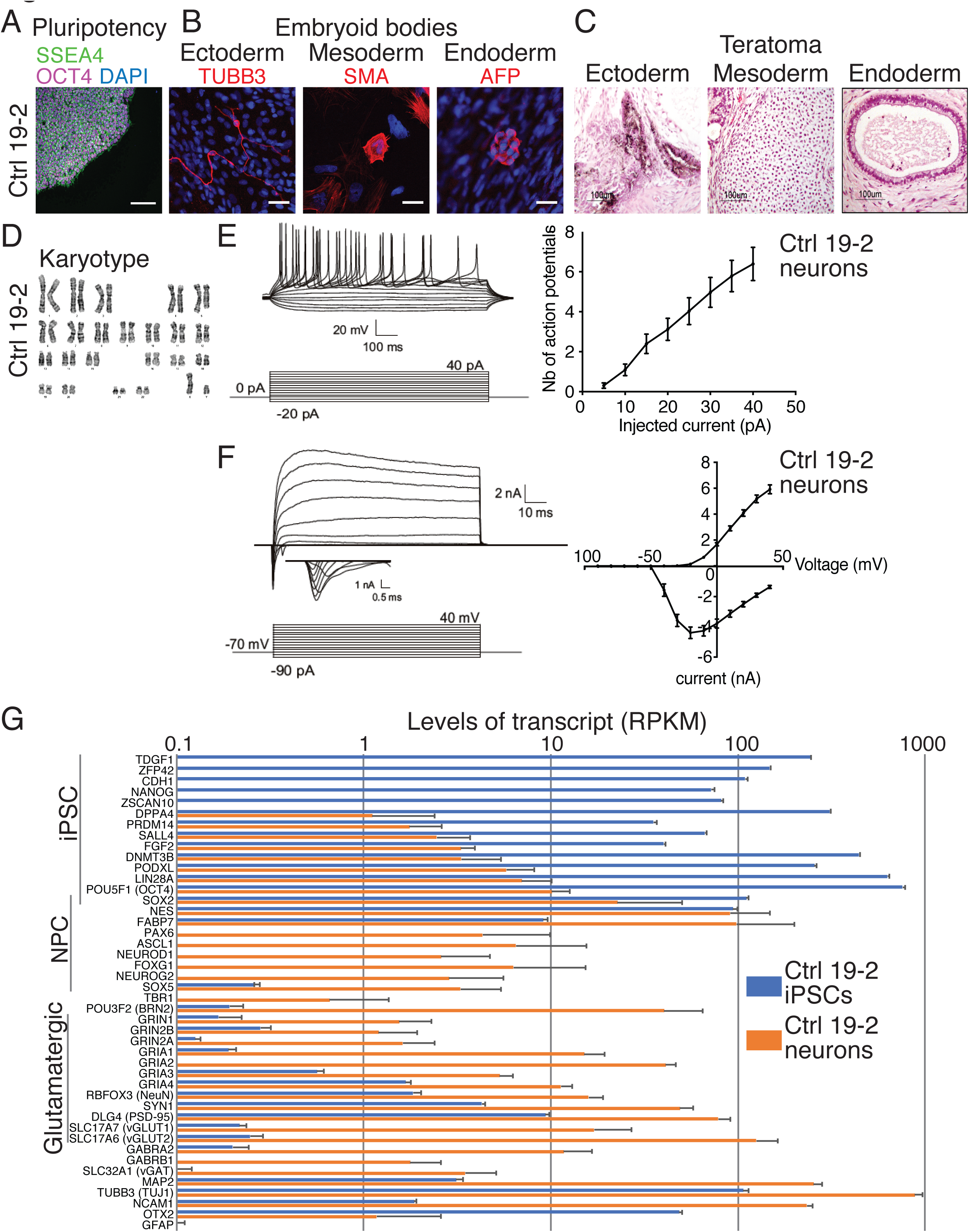
Characterization of the control 19-2 iPSCs and neurons. Representative microscopic images show normal iPSC (A) pluripotency (SSEA4 and OCT4), (B) differentiation potential into the three germ layers in vitro (embryoid bodies: TUBB3, SMA and AFP) or (C) in vivo (teratoma assays), and (D) karyotype. Scale bars: 100μm (A), 25μm (B), and 100μm (C). (E) Representative traces of action potentials recorded at different current injection on the left panel, and the number of action potentials is plotted for each step of current injection on the right panel; (F) Representative traces of sodium currents on the left panel, and currents were recorded at different potentials in voltage-clamp on the right panel; 33 control 19-2 neurons were recorded from three independent differentiation experiments, at day 21-28 post-NEUROG2-induction (PNI). (G) Transcript levels in RPKM of a series of iPSC, neural progenitor cell (NPC) and neuron markers in control 19-2 iPSCs (blue) and control 19-2 glutamatergic neurons (orange). Values are presented as mean±SD of 8 independent experiments for iPSCs and 4 for neurons. mV = millivolt; ms = millisecond; pA = picoampere; nA = nanoampere; μm = micrometer

### Efficiency of Targeting

For 10 of 14 genes, the proper targeting event was achieved (**Table S2**; *ANKRD11, AUTS2, CAPRIN1, CNTNAP2* failed). We did not attempt additional pairs of gRNA. These four unsuccessfully-targeted genes were not necessarily less transcriptionally active than successful genes in iPSCs, as revealed by RNAseq (**Table S2**). For example, *ANKRD11* and *CAPRIN1* presented at reads per kilobase per million mapped reads (RPKM) >25, while *DLGAP2* and *TENM1* were successfully targeted even though at RPKM 0.2 (**Table S2**). A 100% StopTag-VIC signal could come from biallelic StopTag insertion, or from monoallelic StopTag insertion where some indels have disrupted the wt probe sequence on the wt allele. To discern these possibilities, the ratio of wt-FAM:StopTag-VIC alleles was quantified by ddPCR using various combinations of primer/probe sets within the same reaction mix. For example, the amount of StopTag alleles into the *SCN2A* target locus was equivalent to the amount of wt alleles of another autosomal gene, i.e., *ASTN2* (**Figure S2**). Moreover, this amount was equivalent to the double of wt alleles of the allosomal gene *ATRX* (**Figure S2**). These observations support homozygosity of StopTag insertion in *SCN2A*. In addition, we confirmed homozygous, or hemizygous, splicing of StopTag into all successful target loci using PCR amplification and Sanger sequencing, except for *KCNQ2* where the second allele presented a 31-bp frameshift duplication (**Figure 1B**).

### Specificity of Targeting

Each StopTag insertion event was found exclusively on-target as revealed by WGS (**Table S3**). Since the 19-2 genome differs from the hg19 reference genome by probably millions of single nucleotide variants (SNVs), potential extra off-target sites might be missed if we considered only those predicted on hg19. Hence, we have listed all SNV/indel calls unique to each KO line, with respect to the 19-2 control line. Then, we mapped the corresponding gRNA sequences to 200 bp on each side of these calls. We considered as CRISPR-derived on- and off-target variants (**Table S3**) only those with a flanked matching gRNA sequence, tolerating up to five mismatches (**Table S3**). One on-target variant was identified for each of the two gRNAs per gene, corresponding to the expected StopTag insertion (**Table S3**). Remarkably, only one potential off-target mutation was detected within 200 bp of a sequence that had two mismatches with *ATRX* gRNA- (see row 3 in **Table S3**). This single mutation corresponds to a 1-bp deletion located in intron 4 of *IPTK1*, 98 bp away from the Cas9 cut site. Thus, we considered this deletion unlikely to be an off-target effect of CRISPR. It may have arisen as a random spontaneous mutation. Consistent with other findings of high Cas9D10A fidelity (Ran *et al*, 2013; Shen *et al*; 2014), no potential off-target mutation was found within any 200-bp window containing each pair of gRNAs. No off-target mutation was found in any 19-2-derived KO iPSCs (**Table S3**).

### Differentiation into Cortical Excitatory Neurons

Glutamatergic neurotransmission was previously presented as a core pathway associated with ASD (Gilman et al., 2011). Therefore, we sought to differentiate our KO iPSC lines into excitatory neurons to explore functional differences. We used the ectopic expression of NEUROG2 for iPSC differentiation (**Figure 1A**) in order to achieve highly homogeneous populations of neurons for functional testing, and a reproducible differentiation protocol for the simultaneous assessment of multiple cell lines affected by different mutations. Using this protocol, induced neurons, when co-cultured with glial cells, display repetitive action potentials, large inward currents and spontaneous synaptic activity by 21 days in culture (Zhang et al., 2013). We hypothesized that NEUROG2-induced mutant neurons could be used to monitor ASD state *in vitro*. We therefore induced NEUROG2 for 7 days and assessed electrophysiological properties of 19-2 control neurons using patch-clamp recordings of 21-28 days post-NEUROG2-induction (PNI) in presence of cultured mouse glial cells. These 19-2 control neurons had a similar ability to fire repetitive action potentials (**Figure 2E**), and comparable input resistance, inward (sodium) and outward (potassium) currents (**Figure 2F** and **Figure S5A**), as previously reported (Yi et al., 2016; Zhang et al., 2013). Therefore, the NEUROG2 induction protocol was able to generate neurons to a similar maturation as previously reported, enabling us to use the system to interrogate the function of several ASD genes using isogenic knockout iPSCs. Herein, “neuron” will refer to the NEUROG2-induced neuron.

### Transcriptional Characterization of Control 19-2 iPSCs and Neurons

We used RNAseq to verify the pluripotent state of the control 19-2 iPSCs, as well as the glutamatergic state of the control 19-2 neurons four weeks PNI cultured in absence of glial cells. Transcript levels of 14 well-established pluripotency markers were high, i.e., from 33 to 783 RPKM in control iPSCs (**Figure 2G**). Expression of the same set of pluripotency markers was low in neurons, i.e., from 0 to 14 RPKM. We observed a similar pattern of expression to that published originally in NEUROG2-generated neurons (Zhang et al., 2013). For example, we recorded higher levels of the cortical markers *POU3F2* and *FOXG1* than those of *TBR1* (**Figure 2G**). AMPA receptor subunits *GRIA1*, *GRIA2* and *GRIA4* were highly represented at >10 RPKM, compared with the NMDA receptor subunit *GRIN1* at 1.5 RPKM (**Figure 2G**). Transcript levels for the glutamate transporter *SLC17A6* were high, while those of *SLC17A7* were much lower (**Figure 2G**). GABA receptor subunit *GABRA2* transcript levels were seen at higher levels compared with those of the GABA transporter *SLC32A1* (**Figure 2G**). We also observed high levels of the neuronal markers *MAP2*, *TUBB3* and *NCAM1*, but near-zero levels of the astrocyte marker *GFAP* (**Figure 2G**). These data reflect a homogeneous NEUROG2-derived glutamatergic neuron profile, which is suitable for phenotyping assays.

### StopTag Leads to Complete KO of Target Genes Throughout Differentiation

We ensured that the StopTag sequence was properly transcribed and fused to the different target transcripts. For example, a part of some reads generated from RNAseq did not align with exon 2 of *SCN2A* on the human reference genome hg19 (**Figure 3A**), corresponding exactly to the StopTag sequence, which should cause a premature termination of translation. We observed a significant reduction in transcript levels of five target genes in their corresponding KO line (**Figure 3B**). For instance, transcript levels of *ATRX* were reduced by ∼50% in both *ATRX*^−/y^ iPSCs and neurons compared with control cells (**Figure 3B**). This suggests that some mutant transcripts were eliminated by NMD before complete translation due to the presence of a PTC, brought by StopTag. However, we did not observe such reduction in transcript levels for the synaptic genes *CACNA1C*, *KCNQ2*, *SCN2A*, *DLGAP2* and *TENM1* (**Figure 3B** and **Table S4**), suggesting that these transcripts may escape NMD at least until the time of RNA extraction. Despite transcript levels, western blot analysis confirmed the absence of target proteins in mutant neurons. For example, the major protein form of *ATRX* was detected at 280 KDa in control 19-2 neurons, but was not detectable in *ATRX*^−/y^ (**Figure 3C**). The same observation was made for *SCN2A*, at 228 KDa (**Figure 3C**), despite no evident reduction in transcript levels (**Figure 3B** and **Table S4)**.

**Figure 3.**
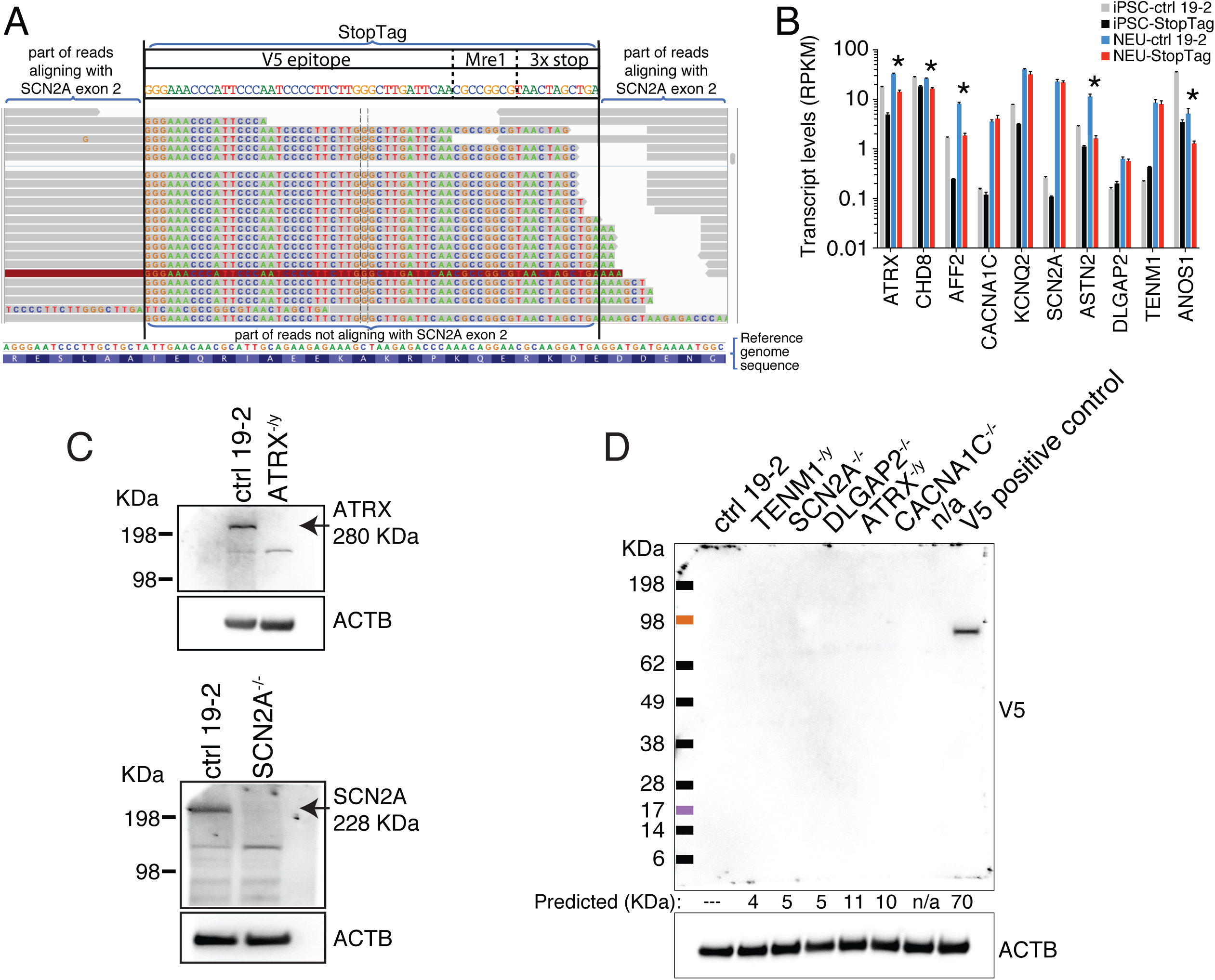
Complete KO of target gene expression in neurons. (A) Example of a target locus, i.e., exon 2 of *SCN2A*, where Spliced Transcripts Alignment to a Reference (STAR) software was used to align reads previously unmapped by TopHat. The grey part of the reads mapped to the human reference genome hg19. The colored part did not map to hg19 but aligned perfectly with the StopTag sequence, showing it is properly transcribed and fused to the target transcripts. (B) Transcript levels in reads per kilobase per million mapped reads (RPKM; y-axis) of each target gene (x-axis) in their corresponding KO iPSCs (black bars) and KO neurons (red bars), as well as in control iPSCs (grey bars) and control neurons (blue bars). Values are presented as mean±SD of 2-8 independent experiments; * = FDR < 0.05 in neurons. (C) Western blots showing the absence of the major form of ATRX (upper panel) and SCN2A (bottom panel) proteins in their corresponding KO neurons, as compared with control 19-2 neurons, four weeks PNI; ACTB: loading control. (D) Western blots revealing the absence of any truncated form of proteins in different KO neuron lines, using a V5 antibody; Predicted (KDa): predicted size of potentially truncated peptides based on the insertion sites of the StopTag within each target transcript relative to the start codon position; n/a: not applicable; ACTB and TUBB3: loading controls. n/a = not available

Notwithstanding, some StopTag-transcripts escaping NMD might still be translated as truncated proteins, and not recognized by antibodies. Such peptides are undesirable in a KO set up since they can present some residual activity or cause other unintended damage (Kamiya et al., 2004; Luo et al., 2017). In order to reveal the presence of any truncated form of proteins, we flanked a sequence coding for a V5 epitope upstream of the 3x stop within the StopTag fragment (**Figure 1A**). Importantly, the SSTR-based splicing of the StopTag sequence was designed to place this V5 within the original translation reading frame. A perfect assemblage was confirmed by Sanger sequencing for most target genes (**Figure 1B**). No truncated forms of protein were detectable by western blot using a V5 antibody (**Figure 3D**), indicating complete absence of target proteins in the testable KO neurons.

### Transcriptional Characterization of KO iPSCs and Neurons

RNAseq profiling of the 10 KO iPSC lines demonstrated their pluripotency and differentiation into cortical neurons. Major iPSC markers, e.g., *NANOG* and *POU5F1*, were highly expressed exclusively in iPSCs (**Figure S3**). Alternatively, specific neuronal markers, e.g., *MAP2* and *SLC17A6* (vGLUT2) were found expressed only in neurons (**Figure S3**). Deficient *GFAP* expression indicated the absence of glial cells in our neuronal cultures (**Figure S3**).

### RNAseq and Pathway Analysis

Transcriptional co-profiling of several isogenic KO lines can potentially reveal how different genes, belonging to different functional groupings, might regulate the expression of common gene-sets or pathways associated with ASD. We explored common pathway enrichment shared by at least three out of the 10 KO lines. Several different gene ontology terms and pathways associated with “Neuron projection development” presented a similar profile in different KO iPSC lines (**Figure 4A**). For example, most of the corresponding gene-sets were down-regulated in *ATRX*-, *ASTN2*- and *DLGAP2*-, while they were upregulated in *AFF2*-null iPSCs (**Figure 4A**). A different profile was observed with “Anchored component of membrane” pathway, i.e., predominantly up-regulated in *AFF2*- and *SCN2A*-, while down-regulated in *ATRX*- and *ASTN2*-null lines (**Figure 4A**). Another group of gene-sets, associated with “Negative regulation of transcription” were commonly down-regulated in *ATRX*-, and up-regulated in *AFF2*- and *SCN2A*-null iPSCs (**Figure 4A**). In general, KO of *CACNA1C* or *KCNQ2* did not have any significant impact on transcriptional networks in iPSCs (**Figure 4A**). This suggests that some of our KO iPSC lines already show transcriptional networks that are prone to ASD-related changes.

**Figure 4.**
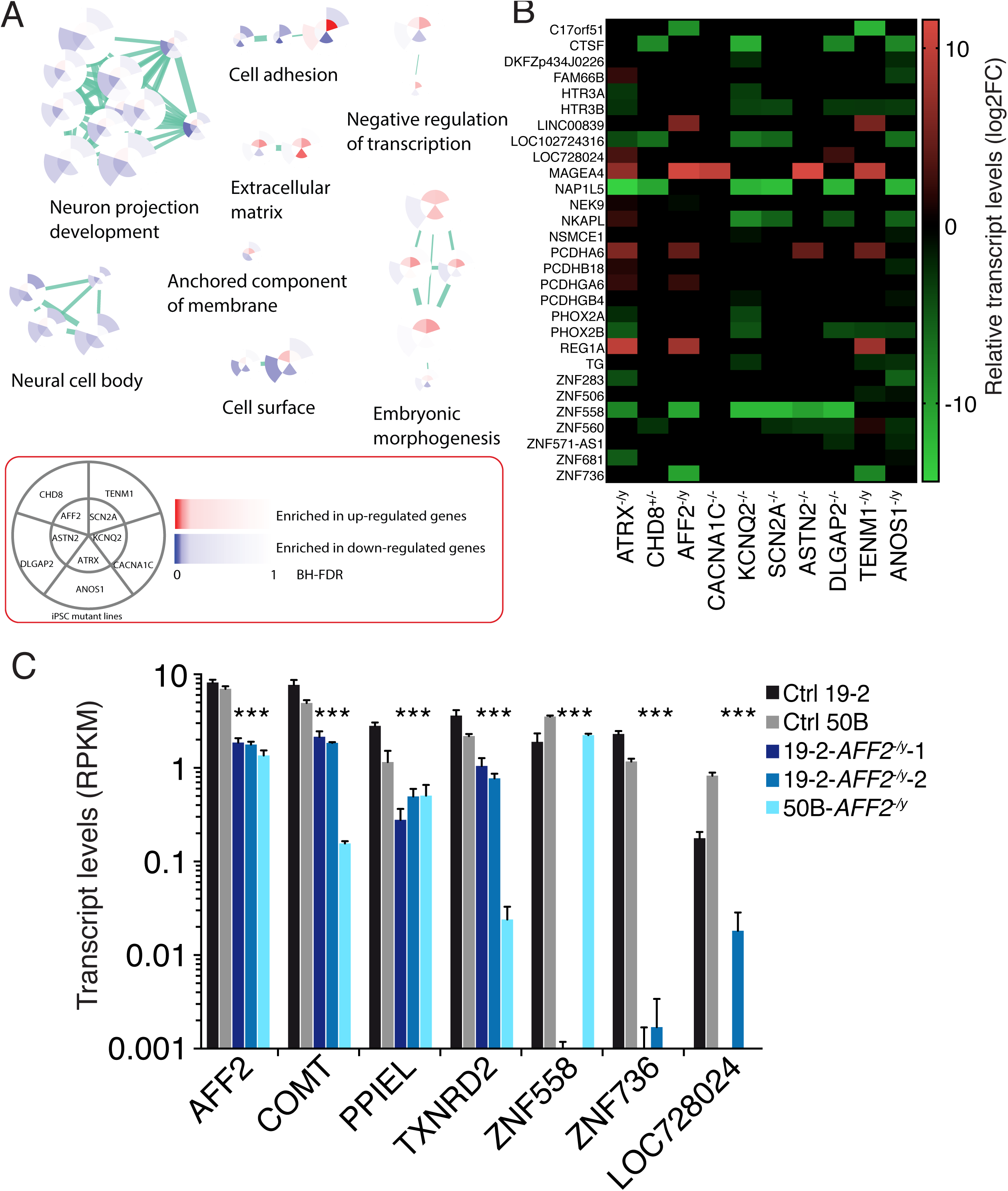
(A) Enrichment map of differentially expressed genes in different mutant iPSCs compared with the isogenic control 19-2, as revealed by RNAseq and pathway analysis. For example, the upper right piece of each pie represents the *SCN2A*-null iPSC line (see asset), in which different gene-sets associated with “Embryonic morphogenesis” are upregulated (red color), with respect to the control 19-2 line. The color intensity correlates with the Benjamini-Hochberg false discovery rate (BH-FDR) values, as depicted in the asset. Values were calculated from (n) independent experiments; n = 8 for ctrl 19-2, n = 4 for *ATRX*^−/y^, *KCNQ2*^−/-^, *SCN2A*^−/-^ and *ASTN2*^−/-^; n = 2 for *AFF2*^−/y^. (B) Differentially expressed genes in different KO neurons compared with the isogenic control 19-2 as revealed by RNAseq. Values are presented as log2 fold change, and only DEGs that passed FDR < 0.05 in at least two different KO lines are shown. Values were calculated from (n) independent differentiation experiments; n = 5 for *AFF2^−/y^*; n = 4 for ctrl 19-2, *ATRX^−/y^*, *KCNQ2*^−/-^, *SCN2A*^−/-^; n = 3 for *ASTN2^−/-^*. (C) RNAseq validation of transcript levels of seven genes in three different *AFF2*-null neuron lines, i.e., 19- 2-*AFF2^−/y^*- 1, 19-2 -*AFF2^−/y^*- 2, and 50B -*AFF2^−/y^*, compared with their respective control lines 19-2 and 50B. Values are presented as mean±SD of (n) independent differentiation experiments where n = 4 for controls 19-2 and 50B; n = 5 for 19-2-AFF2^−/y^-1; n = 3 for 19-2-AFF2^−/y^-2 and 50B-AFF2^−/y^; *each KO line value has a FDR < 0.05 compared to its respective control line.

We also mined RNAseq data for common mechanisms in our iPSC-derived NEUROG2-neurons. However, the number of DEGs passing FDR < 0.05 was substantially lower in induced neurons overall (Table S5) compared to iPSCs (Table S6). No gene ontology terms could significantly be associated with DEGs in most KO neuron lines. This may suggest that the impact on transcriptional networks occurs prior to terminal differentiation of NEUROG2-neurons, e.g., on proliferation of neural progenitor cells (NPCs) as previously shown with *Chd8* in mice (Durak et al., 2016). Therefore, we searched for common DEGs, instead of gene-sets, found in at least two out of ten mutant neuron lines. Several DEGs were shared between distinct KO neuron lines. For instance, *ZNF558* was significantly down-regulated in *ATRX-*, *AFF2-*, *KCNQ2-*, *SCN2A-*, *ASTN2-* and *DLGAP2*-null neurons (**Figure 4B**). Conversely, *REG1A* was up-regulated in *ATRX*-, *AFF2*- and *TENM1*-null neurons (**Figure 4B**). Like in iPSCs, KO of *CACNA1C* did not have a major impact on the neuronal transcriptional network (**Figure 4B**). Interestingly, many up-regulated DEGs shown in **Figure 4B** are also members of the cadherin superfamily (PCDH) involved in synapse configuration (Hirano and Takeichi, 2012), i.e., *PCDHA6*, *PCDHB18, PCDHGA6*, PCDHGB4. Inversely, several down-regulated DEGs belong to the C_2_H_2_ zinc finger superfamily (ZNF) of putative transcription factors (Liu et al., 2014), i.e., *ZNF283, ZNF506, ZNF558, ZNF560, ZNF571-AS1, ZNF681* and *ZNF736* (**Figure 4B**).

Overall, the differential expression analyses in both iPSCs and neurons suggest common pathways and DEGs that converge towards different synaptic factors potentially affecting functional activity of neurons (**
Figure 4A-B**).

### RNAseq Validation in a Different Genetic Background

Isogenic cells are instrumental to control different genetic background contributors. To support our findings, we also suppressed the expression of *AFF2* in a different and unrelated iPSC line. This new line, namely “50B”, was reprogrammed from an unaffected brother of a child with ASD, who was carrying a splice site mutation (c112+1G>C) in the gene *SET* (Yuen et al., 2017). Line 50B does not carry this mutation and was reprogrammed using non-integrative Sendai virus in order to minimize any artefactual genetic modification. This iPSC line presented a normal pluripotency and karyotype (**Figure S4**). We used ribonucleoprotein (RNP) complex as a vector to deliver the CRISPR machinery in 50B iPSCs instead of plasmids. We also used only one gRNA with the Cas9 nuclease for each target gene. We ensured that the sequence of the newly designed gRNA (**Table S1**) was different from that of the gRNAs used in the primary screen, further dissociating any potential off-target mutation to the observed phenotypes.

In addition to this new KO line, i.e., 50B-*AFF2^−/y^*, we generated a second KO line in the 19-2 background using the same method as the first 19-2-*AFF2^−/y^* line. For optimal homogeneity of neuronal cultures, we used the same NEUROG2 differentiation protocol to obtain iPSC-derived glutamatergic neurons, on which we performed RNAseq and differential expression analysis. We searched for significant DEGs in common with all these three *AFF2*-null lines, i.e., 19-2-*AFF2^−/y^*-1, 19-2-*AFF2^−/y^*-2, and 50B-*AFF2^−/y^*, compared with their respective control lines 19-2 and 50B. At least seven DEGs met these criteria, i.e., *AFF2*, *COMT*, *PPIEL*, *TXNRD2*, *ZNF558*, *ZNF736* and *LOC728024* (**Figure 4C**). These results confirm that at least these genes show transcript levels that are influenced by *AFF2*, and not only by the specific 19-2 genetic background.

### Patch-Clamp Electrophysiological Analysis of KO Neurons

To determine if neuronal behaviours were affected by any specific gene KO, we performed patch-clamp recordings on all isogenic KO neuron lines 21 to 28 days PNI. We first measured intrinsic properties of control and KO neuron lines. As shown in **Figure S5A**, overall we did not detect consistent changes in baseline properties in neurons, indicating a similar level of maturity for all KO neurons compared with controls. For example, the capacitance was similar across all lines except for *ATRX*- and *SCN2A*-null neurons where it was lower (**Figure S5A**). The action potential amplitude was unchanged in all lines except for a small reduction in *SCN2A*-null line (**Figure S5A**). While the action potential threshold was reduced in some lines, it didn’t affect the intrinsic firing rate of any of the lines (**Figure S5A**). The only intrinsic property to change across multiple lines was membrane potential, which was elevated in seven of ten lines (**Figure S5A**). This latter finding could either be unrelated to each of the lines, or could have an impact on synaptic activity. Also, it did not correlate with the expression of maturity markers (**Figure S3**). Taken together, these data indicate an overall similar level of maturation for all lines and controls, and individual alterations were not specific to a particular gene. *SCN2A*-deficient neurons were a notable exception since they were significantly altered in all parameters, which is consistent with the known role of sodium currents in regulating neuronal excitability in mouse postnatal neurons (Planells-Cases et al., 2000).

We also measured neuronal activity using patch-clamp electrophysiology as this is perhaps more relevant to ASD. We found that overall, there was a significant reduction in neuronal activity based on the observation of significantly reduced spontaneous EPSC (sEPSC) frequency in *ATRX*-, *AFF2*-, *KCNQ2*-, *SCN2A*- and *ASTN2*-null neurons, without a corresponding change in amplitude (**Figure 5A**). Interestingly, *CACNA1C*^−/-^, *DLGAP2*^−/-^ and *ANOS1*^−/y^ neurons also displayed non-significant reductions in sEPSC frequency (**Figure 5A**). Conversely, *TENM1*^−/y^ tended towards a non-significant higher frequency of sEPSC compared with the isogenic control neurons (**Figure 5A**). Together, these data indicate that ASD risk genes from different classifications can produce a similar electrophysiological phenotype. The common reduction in synaptic activity between these lines indicates that ASD gene networks may converge upon synaptic transmission.

**Figure 5.**
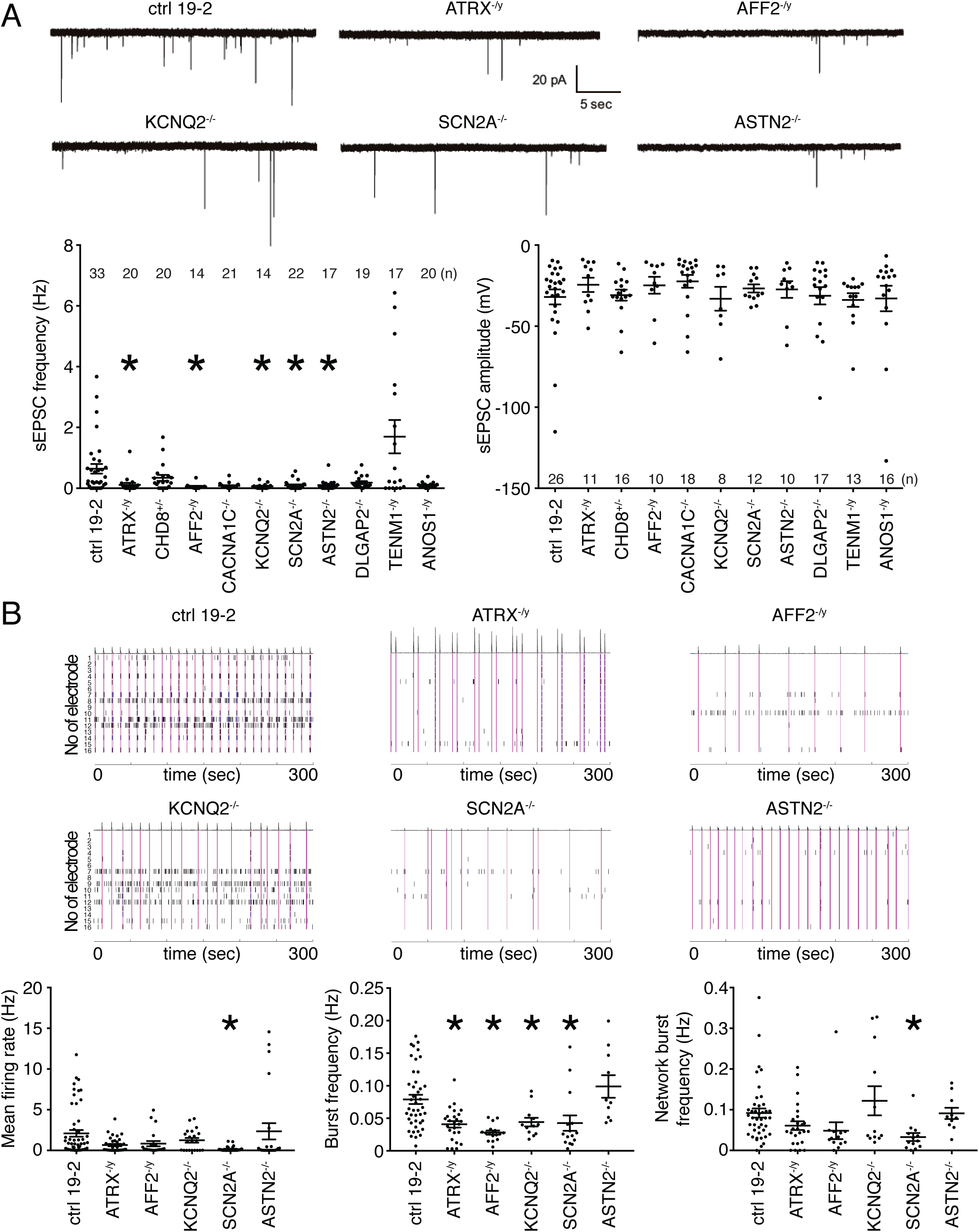
Electrophysiological phenotyping of KO iPSC-derived neurons. (A) Representative traces of excitatory post-synaptic current (EPSC; top panel); spontaneous EPSC frequency (lower left panel) and amplitude (lower right panel) were recorded from different KO neurons; total number of recorded neurons is indicated on the graphs; values are presented as mean±SEM of three independent differentiation experiments for all, except two for *AFF2* and *KCNQ2*, recorded at day 21-28 PNI. (B) Representative raster plots over a 5-minute recording of multi-electrode array experiments (top panels); mean firing rate (lower left panel), burst frequency (lower middle panel), and network burst frequency (lower right panel) were recorded for the five significant genes from the sEPSC frequency graph in (A). Each spike is represented with a short black line. A burst was considered as a group of at least 5 spikes, each separated by an inter-spike interval (ISI) of no more than 100 ms. A network burst (pink lines) was identified as a minimum of 10 spikes with a maximum ISI of 100 ms, occurring on at least 25% of electrodes per well. From 21 to 55 different wells were recorded per line, with usually 6-8 wells per line per experiment. Values are presented as mean±SEM from (n) independent differentiation experiments, where n = 8 for ctrl 19-2; n = 6 for *ATRX*^−/y^; n = 5 for *SCN2A*^−/-^; n = 4 for *AFF2*^−/y^, *ASTN2*^−/-^ and *KCNQ2*^−/-^; recorded at week 8 PNI; * p < 0.05 from one-way ANOVA (Dunnett multiple comparison test). n/d = not detected; pA =picoampere; sec = second; Hz = hertz; mV = millivolt

### Multi-Electrode Array Analysis of KO Neurons

The increased density of neuronal processes > 28 days PNI prevented consistent clean patch-clamp recordings for longer time periods. Nonetheless, a multi-electrode array (MEA) device allows for the ability to monitor the excitability of a population of neurons in an unbiased manner by incorporating the activity of all of individual neurons in a whole-well, and for long time periods in the same culture plate. We monitored the spontaneous neuronal network activity four to eight weeks PNI using MEAs in 48-well format. Field recordings were acquired to estimate the activity in each well. We focused on the five genes that significantly reduced neuronal activity in **Figure 5A**, i.e., *ATRX*, *AFF2*, *KCNQ2*, *SCN2A* and *ASTN2*. We sought to determine if any of these KOs would interfere with synchronized bursting events on a neuronal population level. The highest level of mean firing rate (MFR) was observed eight weeks PNI in control 19-2 neurons (**Figure S5B**). We opted to use this time point for comparison of all KO lines. We also measured the glutamatergic/GABAergic nature of our control 19-2 neurons using different receptor inhibitors. No substantial change was observed after addition of PTX, confirming that GABAergic neurons were not appreciably present in our cultures (**Figure S5C**). However, the MFR was greatly reduced in the presence of CNQX, while unchanged in the non-treated group (group 1 in **Figure S5C**). This suggests that most of the culture was composed of glutamatergic neurons, as expected. Furthermore, all activity was abolished after addition of TTX (**Figure S5C**), indicating the human neurons were expressing functional sodium channels.

The MFR and network burst frequency were significantly reduced in *SCN2A*-null neurons compared to control (**Figure 5B**). Moreover, the burst frequency was lower in *ATRX-, AFF2-*, *KCNQ2-* and *SCN2A-*null neurons (**Figure 5B**). This suggests that loss-of-function of *ATRX, AFF2*, *KCNQ2* or *SCN2A* affects extracellular spontaneous network activity in glutamatergic neurons and reduces neuronal activity on a population level over a longer period of time. Importantly, these data are overall consistent on a per gene level compared with the sEPSC data obtained from individual neurons in **Figure 5A**, except *ASTN2*, and suggest that ASD risk genes converge to disrupt excitatory neuronal activity.

## Discussion

We developed a new CRISPR-based strategy, named StopTag insertion, to completely and specifically knockout the expression of 10 ASD risk genes. This approach allows the generation of multiple isogenic KO lines and pairs well with the highly consistent NEUROG2 induction to study cortical excitatory neurons. This combination is suitable for deep-phenotyping of several isogenic human cortical glutamatergic neuron cultures, which were shown relevant to ASD in several studies (Habela et al., 2016; Moretto et al., 2017). This approach revealed that a common phenotype between diverse ASD risk genes is reduced synaptic activity. We demonstrated that electrophysiological properties of *ATRX*-, *AFF2*-, *KCNQ2*-, *SCN2A*- and *ASTN2*-mutant neurons were severely compromised using two complementary electrophysiology approaches, i.e., patch-clamp and MEA, which allowed for more convincing conclusions when combined together. We also showed that ASD genes from different classes display disruption of common signaling networks that associated with neuron projection and synapse assembly. These results indicate that aberrant functional connectivity is a frequent phenotype in human neurons with ASD candidate gene null mutations.

We obtained a high rate of biallelic editing, i.e., four out of the six successful autosomal genes *CHD8*, *CACNA1C*, *KCNQ2*, *SCN2A*, *ASTN2* and *DLGAP2*. This is possibly due to the use of plasmids to express Cas9 and gRNAs, which are expected to be stable for a few days following nucleofection. We reasoned that the impact of a full KO on transcriptional networks would be more significantly detected compared with heterozygous cells, especially for partially penetrant genes. Many of the candidate genes are located on chromosome X, and since 19-2 cells are of male origin, these genes were fully inactivated after one allele targeted. A notable exception was *CHD8*, which was heterozygous. Despite an attempt to target our *CHD8^+/−^* line in a second round of CRISPR experiment, we failed to isolate any biallelic *CHD8*-null line. It is possible that at least one copy of *CHD8* is indispensable for survival, as proposed previously by other teams regarding mice (Nishiyama et al., 2004) and humans (Bernier et al., 2014). Moreover, the first 39 bp in 5’ of the StopTag sequence was missing in the mutant *CHD8* allele (**Figure 1B**). This left the *Mre1* restriction site and the 3x stop intact. However, the entire V5 epitope was removed. This deletion created a frameshift, now setting the third PTC of the 3x stop within the reading frame instead of the first PTC. The reason for such deletion is unclear. It potentially arose through some form of SSTR-NHEJ co-occurring event.

Different mutations were also found in *KCNQ2*- and *ASTN2*-KO lines, as depicted in **Figure 1B**. For unclear reasons, we could not isolate any full KO line for *ANKRD11*, *AUTS2*, *CAPRIN1* and *CNTNAP2* using this method (grey in **Table S2**). We think that efficiency of the gRNAs used might be responsible, or that targeting these genes might trap the cells with a proliferative disadvantage, being constantly outnumbered by untargeted cells during the enrichment steps.

We believe that the NEUROG2-neuron system represents a substantial advantage compared with classic dual-SMAD inhibition differentiation protocols for phenotyping experiments that require relatively high levels of cell homogeneity. Dual-SMAD inhibition-neurons are more heterogeneous, including several different types of cells, they need a longer time to mature, and may not be ideal for higher-throughput studies where multiple genes need to be compared. We found that scalability and consistency are remarkably improved by the NEUROG2 differentiation protocol. Moreover, it allows to test directly the neuronal phenotype and eliminates any contribution from mutant glial cells. However, one caveat to the NEUROG2 approach is the lack of inhibitory neurons, which may be important to analyze particular ASD risk genes expressed in this population, e.g., *ARID1B* (Jung et al., 2017). In addition, our data suggest that a disruption in the excitation/inhibition balance contributes to ASD (Hussman, 2001; Rubenstein and Merzenich, 2003). However, to test this directly, the effect on inhibitory neurons will have to be evaluated in the future. The recent generation of induced GABAergic neurons (Yang et al., 2017) should allow such analysis of ASD risk genes in inhibitory neurons, as well as in co-cultures of excitatory and inhibitory neurons.

Interestingly, the five genes with a common sEPSC reduced neuronal activity phenotype (**Figure 5A**) fall into different molecular function groupings, i.e., *ATRX* in transcriptional regulation, *AFF2/FMR2* in RNA processing, and *KCNQ2, SCN2A, ASTN2* in synaptic and adhesion (**Table 1**). And KO of three of them, i.e., *ATRX*, *AFF2*, *SCN2A* (from three different groupings), converge to common transcriptional networks associated with, for example, “Transcription regulation”, “Membrane components” and “Embryonic morphogenesis” in iPSCs (**Figure 4A**), suggesting that early developmental molecular events can already trap the embryo on the course of ASD.

We also found common DEGs among the five sEPSC-featured mutant neurons, e.g., *PCDHA6*, *REG1A* and *ZNF558* (**Figure 4B**) that might be responsible for this common phenotype. These results suggest that RNAseq could be used to detect specific transcriptional signatures associated with ASD. Several DEGs were also shared between neurons mutant for genes from the same group, e.g., *KCNQ2* and *SCN2A*, which are both ion channel subunits; and between neurons mutant for *ATRX* and *AFF2*, which are both X-linked genes involved in intellectual disability and binding G4-quadruplexes associated with DNA and RNA, respectively (Bensaid et al., 2009; Law et al., 2010) (**Figure 4B**). This raises the possibility that G4-quadruplexes contribute to the establishment of ASD cellular phenotypes. However, it is also possible that not all DEGs may be contributing to the synaptic phenotype, potentially involving post-translational mechanisms. For example, a recent study identified that CASK protein levels, not mRNA levels, were impacted by a deletion of the ASD and schizophrenia risk gene *NRXN1* (Pak et al., 2015). Future proteomic studies may indeed be needed to identify similar mechanisms in our isogenic neuron lines, as demonstrated with different neurons (Djuric et al., 2017). Regardless, the reduced activity phenotype in our KO cell lines suggests a convergence between different signaling systems in neurons, indicating that pharmacological interventions that correct the synaptic phenotype could be applied to diverse genetic causes of ASD (Lai et al., 2014; Sahin and Sur, 2015). Although the total number of DEGs was relatively low in our KO NEUROG2-neurons, some were validated in a different genetic background (line 50B in **Figure 4C**) and were previously associated with ASD, e.g., *COMT* is involved in the transfer of a methyl group and inactivation of catecholamine neurotransmitters and was associated with schizophrenia (Lee et al., 2005). Furthermore, *PPIEL* is a pseudogene whose DNA methylation level was associated with intellectual disability and bipolar disorder (Kuratomi et al., 2008).

The reason why the other five KO neuron lines, i.e., *CHD8^+/−^*, *CACNA1C^−/-^*, *DLGAP2^−/-^*, *TENM1^−/y^* and *ANOS1^−/y^* did not reach statistical significance in the sEPSC readout remains elusive. *CHD8* has extensively been associated with ASD but, as mentioned above, it seems to influence the proliferation of neural progenitor cells and to a lesser extent the function of post-mitotic cortical glutamatergic neurons (Durak et al., 2016; Platt et al., 2017). *CACNA1C* is responsible for the Timothy syndrome (**Table 1**), which is associated with cardiac abnormalities, but did not present any significant phenotype in this setting, except a different resting membrane potential in **Figure S5A**. *De novo* duplications of *DLGAP2* were found in patients with ASD (Marshall et al., 2008; Pinto et al., 2010). However, DLGAP2 is a synaptic protein associated with NMDA receptors which may be underrepresented in 21-28 days PNI, necessitating further experiments. *ANOS1*, which is responsible for the Kallmann syndrome (**Table 1**) and seems to affect predominantly gonadotropin-releasing neurons (Hardelin et al., 1993), only presented a different resting membrane potential in NEUROG2-neurons ( **Figure S5A)**. A non-sense mutation in *TENM1* was found in a patient with ASD who co-carries a missense mutation in *AFF2* (Yuen et al., 2017), and only a different action potential threshold was detected in *TENM1*-null neurons (**Figure S5A)**. As additional genetic studies are performed, the priority ranking of all of the genes in our study for a role in ASD will likely continue to shift, and in some cases, be influenced by functional studies such as are performed here.

In conclusion, we have designed a precise CRISPR gene editing strategy for complete KO of a series of 10 ASD risk genes from different classifications, and shown that they can be responsible for similar transcriptional rewiring and electrophysiological phenotypes in human iPSC-derived glutamatergic neurons. Overall, given the heterogeneity involved in ASD, we believe that this type of CRISPR-isogenic KO system is essential for properly controlled cellular phenotyping experiments. Whole animal murine KO models would also be useful but the same efficiencies would not be possible. For future experiments, our isogenic KO system could also be used for the incremental creation of isogenic knock-in lines where other ASD patient mutations, combinatorically, could be introduced creating a resource for advanced functional modeling and new therapeutic testing.

## Acknowledgements

We thank the Autism Speaks MSSNG project for genomic data and linking to consented families. We also thank Melissa Carter, Wendy Roberts, Brian Chung and Rosanna Weksberg for obtaining skin biopsies and blood work, and the families for volunteering. We also thank Sergio Pereira, Wilson Sung, Thomas Nalpathamkalam, Darwin D’Souza, Joe Whitney, Jeff MacDonald and Liz Li for bioinformatic support; Tara Paton, Guillermo Casallo, Barbara Kellam, Ny Hoang and Sylvia Lamoureux for technical help; T.C. Südhof for the NEUROG2/rtTA lentiviral constructs.

## Author Contributions

E.D., R.K.C.Y., K.K.S., J.E. and S.W.S designed the research project. E.D., S.H.W., M.F., P.J.R., W.W. and A.P. contributed to cell maintenance, characterization and differentiation. E.D. conceived the StopTag KO strategy. E.D., M.F. and K.Z. contributed to CRISPR experiments. Z.W. performed WGS off-target analyses. D.C.R. performed Western blots. R.A., G.P., B.T., E.D., G.K. and D.M. participated in RNAseq and pathway analysis. E.D. and S.H.W. performed electrophysiological experiments. J.L.H., V.K., S.W. and P.P. provided technical help. E.D., S.H.W., K.K.S., J.E., and S.W.S. wrote the manuscript and supervised the project. Specific contributions of the co-corresponding authors: K.K.S. lab differentiated iPSCs to neurons and conducted patch-clamp electrophysiology experiments; J.E. lab generated iPSC. and differentiated to neurons for expression analyses and multi-electrode arrays; S.W.S. lab performed the CRISPR editing, WGS and RNAseq experiments.

## Data Availability

All RNAseq data was deposited in GEO database under the accession number GSE107878.

## Experimental Procedures

Reprogramming of iPSCs was performed under the approval of the Canadian Institues of Health Research Stem Cell Oversight Committee, and the Research Ethics Board of The Hospital for Sick Children, Toronto.

### Skin Fibroblast Culture

The skin-punch biopsy for the control 19-2 came from the upper back by a clinician at The Hospital for Sick Children, with approval from their Research Ethics Board. Samples were immersed in 14 ml of ice-cold Alpha-MEM (Wisent Bioproducts) supplemented with penicillin 100 units/ml and streptomycin 100 μg/ml (pen/strep; ThermoFisher), and transferred immediately to the laboratory. The biopsy was cut into ∼1mm^3^ pieces in a 60-mm dish. 5 ml of collagenase 1 mg/ml (Sigma) was added and the dish was placed in 37°C incubator for 1:45 hours. Skin pieces and collagenase were then transferred to a 15-ml tube, and centrifuged at 300 g for 10 minutes. Supernatant was removed and 5 ml of trypsin 0.05%/EDTA 0.53 mM (Wisent Bioproducts) was added. The mix was pipetted up and down several times to break up tissue and placed in 37°C incubator for 30 minutes. After incubation, the mix was centrifuged at 300 g for 10 minutes, and the supernatant was removed, leaving 1 ml. The pellet was pipetted up and down vigorously to break the pieces. The mix was transferred in a T-12.5 flask along with 5 ml of Alpha-MEM, 15% Fetal Bovine Serum (FBS; Wisent Bioproducts), 1x pen/strep, and placed in a 37°C incubator. Cultured cells were fed every 5 days. At 100% confluency, the cell population was split into three 100-mm dishes for expansion and cryopreservation in liquid nitrogen.

### Skin Fibroblast Reprogramming

The iPSC line 19-2 was generated from skin fibroblasts using retroviral vectors encoding *POU5F1*, *SOX2*, *KLF4*, *MYC*, and lentiviral vectors encoding the pluripotency reporter EOS-GFP/Puro^R^, as described previously (Hotta et al., 2009).

### Peripheral Blood Mononuclear Cells (PBMCs) Isolation and Enrichment of CD34+ Cells

Whole peripheral blood was processed at the Centre for Commercialization of Regenerative Medicine (CCRM) using Lymphoprep™ (StemCellTechnologies) in a SepMate™ tube (StemCellTechnologies) according to manufacturer’s instructions. The sample was centrifuged 10 min at 1200 g. The top layer containing PBMCs was collected and mixed with 10 ml of the PBS/FBS mixture and centrifuged 8 min at 300 g. The PBMC’s collected at the bottom of the tube were washed, counted and re-suspended in PBS/FBS mixture. CD34+ cells were isolated using the Human Whole Blood/Buffy Coat CD34+ Selection kit according to manufacturer’s instructions (StemCellTechnologies). Isolated cells were expanded in StemSpan SFEM II media (StemCellTechnologies) and StemSpan CD34+ Expansion Supplements (StemCellTechnologies) prior to reprogramming.

### Reprogramming PBMC Using Non-Integrative Sendai Virus

The iPSC line 50B was generated from CD34+ PBMCs at CCRM using CytoTune ™-iPSC 2.0 Sendai Reprogramming Kit. Expanded cells were spun down and resuspended in StemSpan SFEM II media and StemSpan CD34+ Expansion Supplements, and placed in a single well of a 24-well dish. Virus MOI was calculated and viruses combined according to number of cells available for reprogramming and manufacturer’s protocol. The virus mixture was added to cells, and washed off 24 hours after infection. 48 hours after viral delivery, cells were plated in 6-well plates in SFII and transitioned to ReproTESR for the duration of reprogramming. Once colonies were of an adequate size and morphology, individual colonies were picked and plated into E8 media (ThermoFisher). Clones were further expanded and characterized using standard assays for pluripotency, karyotyping (Medical Genetics Services, Addenbrooke’s Hospital), short-tandem repeat (STR) genotyping [The Centre for Applied Genomics (TCAG)] and mycoplasma (Lonza MycoAlert). Directed differentiation was performed using kits for definitive endoderm, neural and cardiac lineages (ThermoFisher).

### iPSCs Maintenance

All iPSC lines were maintained on matrigel (Corning) coating, with complete media change every day in mTeSR™ (StemCellTechnologies). ReLeSR™ (StemCellTechnologies) was used for passaging. Accutase™ (InnovativeCellTechnologies) and 10 μM Rho-associated kinase (ROCK) inhibitor (Y-27632; StemCellTechnologies) were used for single-cell dissociation purposes.

### Gene Editing

Guide RNA sequences were devised using the CRISPR design tools (Hsu et al., 2013) or (http://www.benchling.com) in order to minimize off-target activity. We selected the highest off-target and on-target scored pair of gRNAs with minimal inclusion of single nucleotide variants with respect to the control 19-2 genome. Two different methods were used to prepare the gRNAs. The first method involved cloning of gRNAs into expression plasmids. DNA sequences allowing fusion of crRNAs and tracrRNAs were designed and assembled as described (Yang et al., 2014) and were synthesized at TCAG. The assembled inserts were ligated into plasmid #41824 (Addgene) using Gibson Assembly^®^ cloning kit [New England Biolabs (NEB)]. HEK293T cells (ATCC) were transfected, as 200 000 cells per well in 24-well plate, with 200 ng of each gRNA plasmids, 400 ng of pCas9D10A_GFP plasmid #44720 (Addgene), and 1 μl of 100 μM ssODN synthesized as standard desalted Ultramer^®^ oligonucleotides [Integrated DNA Technologies (IDT)], using lipofectamine^®^ 2000 (ThermoFisher). For human iPSCs, 3 μg of each gRNA plasmids, 5 μg of pCas9D10A_GFP plasmid, and 3 μl of 100 μM ssODN were nucleofected into 1×10^6^ iPSCs using Nucleofector ™2b device (Lonza; program A-023) and Human Stem Cell Nucleofector^®^ kit 1 (Lonza). The second method to prepare gRNAs involved ribonucleoprotein (RNP) complexes. For this, crRNA and tracrRNA were synthesized and annealed according to manufacturer’s protocol (Synthego). 1.6 μl of 10 μM gRNA was incubated with 1 μl of 20 μM Cas9-2NLS Nuclease (Synthego) in 100 μl Human Stem Cell Nucleofection Solution 1 (Lonza) for 10 minutes at room temperature to form Cas9:gRNA RNP complexes. Then, 5 μl of 10 μM ssODN was added to the complexes. The mix was nucleofected into 1×10^6^ iPSCs using Nucleofector™2b (program A-023). Prior to nucleofection, iPSCs were treated with 10 μM Y-27632 for 60 minutes at 37°C. Following nucleofection, iPSCs were evenly distributed into a flat-bottom 96-well plate in mTeSR and 10 μM Y-27632.

### Purification of Edited Cells

The method used to enrich for desired edited iPSCs was based on droplet digital PCR (ddPCR) and enrichment steps, and was adapted from Miyaoka et al. (Miyaoka et al., 2014). Near-confluent iPSCs, i.e., 10-14 days post-nucleofection or post-passaging, were treated with 10 μM Y-27632 for 60 minutes at 37°C, washed with PBS, and treated with 28 μl/well of accutase for 10 minutes at 37°C. Half of each well was harvested in a PCR 96-well plate containing 50 μl/well of DNA lysis buffer (10mM Tris pH 7.5, 10mM EDTA pH 8.0, 10mM NaCl, 0.5% Nlauroylsarcosine and freshly added 1 mg/ml proteinase K). The other half was reseeded in a new flat-bottom matrigel-coated 96-well plate along with 250 μl/well of mTeSR™ supplemented with 10 μM Y-27632, and put at 37°C for expansion. The PCR plate containing cells and DNA lysis buffer was sealed and heated at 70°C for 10 minutes in a PCR machine, and chilled on ice for 3 minutes. 100ul/well of DNA precipitation solution (EtOH 95%, H_2_O 5%, NaCl 75mM), previously cooled at −80°, was added and the plate was left at room temperature for 30-60 minutes. The plate was spun at 1600 g for 4 minutes and flicked upside down quickly to remove supernatant. DNA was washed 2 consecutive times with 100ul/well EtOH 70%, and heated without seal at 70°C for 2 minutes. DNA was resuspended with 30ul/well of TE buffer, the plate was sealed and heated at 70°C for 10 minutes. The ddPCR (Bio-Rad) was performed according to manufacturer’s protocol. Custom TaqMan^®^ MGB probes (ThermoFisher) were designed according to manufacturer’s recommendations. A DNA probe specific to StopTag, which encompassed an *Mre*1 restriction site and the first half of the 3x stop of StopTag, was coupled with the fluorescent dye VIC (**Table S1**). This design ensured high specificity of the StopTag-VIC probe and secured the isolation of cells that have integrated the 3x stop at the target locus. Conversely, a different probe coupled with the fluorescent dye FAM was used for each unmodified (wt) target gene. Each wt-FAM probe sequence was selected to overlap with that of reverse gRNA (gRNA-; **Table S1**), which instructs Cas9D10A to nick the plus strand. The entire wt-FAM probe sequence was removed from ssODN to avoid confusion with StopTag allele detection. The Tm of the common VIC probe was ensured to be compatible with the Tm of all other specific FAM probes. Moreover, about half of the sequence of gRNA- was removed from ssODN to prevent nicking from Cas9D10A, given that ssODN was synthesized as plus strand for each target gene. The forward primers for ddPCR were designed within the ssODN, and reverse primers outside of ssODN, to prevent PCR amplification of non-spliced ssODN sequences (**Table S1**). Primers were designed using primer blast tool (www.ncbi.nlm.nih.gov/tools/primer-blast/). The well with the highest frequency of edited alleles was identified in absolute quantification mode. When the re-seeded half of cells reached confluency, 1/10 of total cell number of the corresponding selected well were passed in a new flat-bottom matrigel-coated 96-well plate in order to enrich for edited cells while maintaining a polyclonal composition of cell population. When confluent, this secondary plate was processed for ddPCR, as described above. This procedure was repeated until a 100% edited iPSC population was obtained. For characterization of the new 100% edited lines, a PCR was performed on genomic DNA from KO cells using OneTaq^®^ DNA polymerase (NEB) according to manufacturer’s protocol in order to amplify a 500-1000 fragment encompassing the StopTag integration site. Amplified fragments were ligated into plasmids using TOPO TA cloning kit (ThermoFisher). These plasmids were Sanger sequenced to reveal the exact sequence and zygosity of each target locus.

### Detection of CRISPR-Related Off-Target Genetic Variants

Genomic DNA was extracted from 70-80% confluent iPSCs using DNeasy^®^ tissue kit (Qiagen), and submitted to TCAG for genomic library preparation and whole genome sequencing (WGS). DNA samples were quantified and analyzed using Qubit High Sensitivity Assay and Nanodrop OD260/280 ratio. 200 ng of DNA was used as input material for library preparation using Illumina TruSeq Nano DNA Library Prep Kit. DNA was fragmented to 550 bp on average using sonication on a Covaris LE220 instrument. Fragmented DNA was end-repaired and A-tailed. Indexed TruSeq Illumina adapters with overhang-T were added to the DNA. Libraries were loaded on a Bioanalyzer DNA High Sensitivity chip to validate size and absence of primer dimers, and were quantified by qPCR using Kapa Library Quantification Illumina/ABI Prism Kit protocol (KAPA Biosystems). Each validated library was sequenced on two lanes of a high throughput V4 flowcell on a HiSeq 2500 platform following Illumina’s recommended protocol to generate paired-end reads of 125-bases in length. These reads were mapped using the bwa algorithm (0.7.12). GATK (GenotypeConcordance) (McKenna et al., 2010) was used to identify indel and SNV calls unique to the different KO line genomes with respect to the control 19-2 genome. Because of the 59-bp length of the StopTag sequence, it could have appeared as a series of GATK calls in close proximity to one another. Therefore, all calls occurring within a 200 bp window were grouped, and GATK FastaAlternateReferenceMaker was used to reconstruct the sequence within that window to identify any potential insertion. We found StopTag insertions only at the expected loci. Then, we searched for sequences matching (up to five mismatches) gRNA+ and gRNA-, separately, within a 200-bp window flanking both sides of each unique variant call, in each corresponding KO line. Since the gRNA cloning protocol that we used sets the first 5’ nucleotide as G, which is dispensable for specificity (Yang et al., 2014), mismatches at that position were ignored. Likewise, mismatches at the PAM sequence were ignored.

### Lentivirus Production

7.5×10^6^ HEK293T cells were seeded in a T-75 flask, grown in 10% fetal bovine serum in DMEM (Gibco). The next day, cells were transfected using Lipofectamine 2000 with plasmids for gag-pol (10 μg), rev (10 μg), VSV-G (5 μg), and the target constructs FUW-TetO-Ng2-P2A-EGFP-T2A-puromycin or FUWrtTA (15 μg; gift from T.C. Südhof laboratory) (Zhang et al., 2013). Next day, the media was changed. The day after that, the media was spun down in a highspeed centrifuge at 30,000 g at 4°C for 2 hours. The supernatant was discarded and 50 μl PBS was added to the pellet and left overnight at 4°C. The next day, the solution was triturated, aliquoted and frozen at −80°C.

### Differentiation into Glutamatergic Neurons

5×10^5^ iPSCs/well were seeded in a matrigel-coated 6-well plate in 2 ml of mTeSR supplemented with 10 μM Y-27632. Next day, media in each well was replaced with 2 ml fresh media plus 10 μM Y-27632, 0.8 μg/ml polybrene (Sigma), and the minimal amount of NEUROG2 and rtTA lentiviruses necessary to generate 100% GFP+ cells upon doxycycline induction, depending on prior titration of a given virus batch. The day after, virus-containing media were replaced with fresh mTeSR, and cells were expanded until near-confluency. Newly generated “NEUROG2-iPSCs” were detached using accutase, and seeded in a new matrigel-coated 6-well plate at a density of 5×10^5^ cells per well in 2 ml of mTeSR supplemented with 10 μM Y-27632 (day 0 of differentiation). Next day (day 1), media in each well was changed for 2 ml of CM1 [DMEM-F12 (Gibco), 1x N2 (Gibco), 1x NEAA (Gibco), 1x pen/strep (Gibco), laminin (1 μg/ml; Sigma), BDNF (10 ng/μl; Peprotech) and GDNF (10 ng/μl; Peprotech) supplemented with fresh doxycycline hyclate (2 μg/ml; Sigma) and 10 μM Y-27632. The day after (day 2), media was replaced with 2 ml of CM2 [Neurobasal media (Gibco), 1x B27 (Gibco), 1x glutamax (Gibco), 1x pen/strep, laminin (1 μg/ml), BDNF (10 ng/μl) and GDNF (10 ng/μl)] supplemented with fresh doxycycline hyclate (2 μg/ml) and puromycin (5 μg/ml for 19-2-derived cells, and 2 μg/ml for 50B-derived cells; Sigma). Media was replaced with CM2 supplemented with fresh doxycycline hyclate (2 μg/ml). The same media change was repeated at day 4. At day 6, media was replaced with CM2 supplemented with fresh doxycycline hyclate (2 μg/ml) and araC (10 μM; Sigma). Two days later, these day 8 post-NEUROG2-induction (PNI) neurons were detached using accutase and ready to seed for subsequent experiments, as described below.

### RNAseq and q-RT-PCR

Total RNA was extracted from 70-80% confluent iPSCs using RNeasy^®^ mini kit (Qiagen). For glutamatergic neurons, 12-well plates were coated with filter-sterilized 0.1% polyethyleneimine (PEI; Sigma) solution in borate buffer pH 8.4 for 1 hour at room temperature, washed four times with water, and dried thoroughly. 1×10^6^ neurons/well (day 8 PNI) were seeded in 2 ml CM2 media. Media was half-changed once a week with CM2 media. Four weeks later, total RNA was extracted from each well using RNeasy^®^ mini kit (Qiagen). RNA samples were submitted to an Agilent Bioanalyzer 2100 RNA Nano chip for quality control. Concentration was determined using Qubit RNA HS Assay on a Qubit fluorometer (ThermoFisher). RNA libraries were prepared using NEBNext Ultra RNA Library Preparation kit for Illumina. Briefly, 500 ng of total RNA was used for poly-A mRNA enrichment before being split into 200-300bp fragments for 4 minutes at 94°C. Fragments were converted to double stranded cDNA, end-repaired and adenylated in 3’ to create an overhang A, allowing ligation of Illumina adapters with an overhang T. Fragments were amplified under the following conditions: initial denaturation at 98°C for 30 seconds, followed by 13 cycles of 98°C for 10 seconds, 65°C for 75 seconds, and finally an extension step of 5 minutes at 65°C. Each sample was amplified with a different barcoded adapter to allow for multiplex sequencing. RNA library integrity was verified on a Bioanalyzer 2100 DNA High Sensitivity chip (Agilent Technologies), and quantified using Kapa Library Quantification Illumina/ABI Prism Kit protocol (KAPA Biosystems). Stranded libraries were pooled in equimolar amounts and sequenced on an Illumina HiSeq 2500 platform using a High Throughput Run Mode flowcell and the V4 sequencing chemistry to generate paired-end reads of 125-bases in length, following Illumina’s recommended protocol. Data quality was assessed using FastQC v.0.11.2. Trimmed reads were screened for presence of rRNA and mtRNA sequences using FastQ-Screen v.0.4.3. RSeQC package v.2.3.7 was used to assess read distribution, positional read duplication and confirm strandedness of alignments. Raw trimmed reads were aligned to the reference genome hg19 using Tophat v.2.0.11. Tophat alignments were processed to extract raw read counts for genes using htseq-count v.0.6.1p2. Two-condition differential expression was done with the edgeR R package, v.3.8.6. In the filtered data set were retained only genes whose reads per kilobase of expressed transcript per million reads of library (RPKM) values were > 1 for the iPSCs and 0.5 for neurons in at least n-1 samples where n is the number of samples in the smaller sample group in the comparison. The method used for normalizing the data was TMM, implemented by the calcNormFactors (y) function. All samples in each pair-wise comparison were normalized together. The test for differential expression was done using the quasi-likelihood F-test functionality in edgeR. For q-RT-PCR, total RNA was reverse transcribed into cDNA using First Strand cDNA Synthesis kit according to manufacturer’s protocol (Sigma), and quantitative PCR was performed using SYBR^®^ Select Master Mix kit (LifeTechnologies) with primer sets designed using primer blast tool.

### Pathway Enrichment

Pathway enrichment analysis was performed using the R package goseq version 1.28.0 and R version 3.4.1 (2017-06-30) using a custom gene-set collection including gene ontology (GO, obtained from the R package GO.db version 3.4.1) and pathways (KEGG and Reactome collections downloaded from the respective websites on 20171016). To remove terms that were either too generic or too narrow, GO size terms were restricted to those within 100-1000 annotated genes, and pathways to 15 and 500 annotated genes, leaving 3736 total gene ontology terms and pathways including 17041 genes. Up- and down-regulated genes were separately analyzed using goseq (Wallenius’ noncentral hypergeometric distribution); median transcript lengths were used to correct for count bias. To correct for multiple comparisons, Benjamini-Hochberg false discovery rate (BH-FDR) correction was applied to the p-values. The results were combined to obtain a list of pathways enriched in up-regulated and down-regulated genes.

### Patch-Clamp Recordings

Day 3 PNI neurons were replated at a density of 100 000/well of a polyornithin/laminin coated coverslips in a 24-well plate with CM2 media. On day 4, 50 000 mouse astrocytes were added to the plates and cultured until day 21-28 PNI for recording. At day 10, CM2 was supplemented with 2.5% FBS in accordance with (Zhang et al., 2013). Whole-cell recordings (BX51WI; Olympus) were performed at room temperature using an Axoclamp 700B amplifier (Molecular Devices) from borosilicate patch electrodes (P-97 puller; Sutter Instruments) containing a potassium-based intracellular solution (in mM): 123 Kgluconate, 10 KCL, 10 HEPES; 1 EGTA, 2 MgCl_2_, 0.1 CaCl_2_, 1 Mg-ATP, and 0.2 Na_4_GTP (pH 7.2). 0.06% sulpharhodamine dye was added to select neurons for visual confirmation of multipolar neurons. Composition of extracellular solution was (in mM): 140 NaCl, 2.5 KCl, 1 1.25 NaH_2_PO_4_, 1 MgCl_2_, 10 glucose, and 2 CaCl_2_ (pH 7.4). Whole cell recordings were clamped at −70 mV using Clampex 10.6 (Molecular Devices), corrected for a calculated −10 mV junction potential and analyzed using the Template Search function from Clampfit 10.6 (Molecular Devices). Following initial breakthrough and current stabilization in voltage clamp, the cell was switched to current clamp to monitor initial spiking activity and record the membrane potential (cc=0, ∼1 min post-breakthrough). Bias current was applied to bring the cell to ∼70 mV whereby increasing 5 pA current steps were applied (starting at −20 pA) to generate the whole cell resistance and to elicit action potentials. Data were digitized at 10 kHz and low-pass filtered at 2 kHz. Inward and outward currents were recorded in whole-cell voltage clamp in response to consecutive 10 mV steps from −90 mV to +40 mV.

### Multi-electrode Array

48-well opaque- or clear-bottom MEA plates (Axion Biosystems) were coated with filter-sterilized 0.1% PEI solution in borate buffer pH 8.4 for 1 hour at room temperature, washed four times with water, and dried overnight. 120 000 day 8 PNI neurons (were Dox-induced for 7 days) per well were seeded in 250 μl CM2 (laminin 10 μg/ml instead of 1 μg/ml for MEA) media. The day after, 10 000 mouse astrocytes/well were seeded on top of neurons in 50 μl/well CM2 media. Astrocytes were prepared from postnatal day 1 CD-1 mice as described (Kim and Magrane, 2011). Media was half-changed once a week with CM2 media. Once a week from week 4 to 8 PNI, the electrical activity of the MEA plates was recorded using the Axion Maestro MEA reader (Axion Biosystems). The heater control was set to warm up the reader at 37°C. Each plate was first incubated for 5 minutes on the pre-warmed reader, then real-time spontaneous neural activity was recorded for 5 minutes using AxIS 2.0 software (Axion Biosystems). A bandpass filter from 200 Hz to 3 kHz was applied. Spikes were detected using a threshold of 6 times the standard deviation of noise signal on electrodes. Offline advanced metrics were re-recorded and analysed using Axion Biosystems Neural Metric Tool. An electrode was considered active if at least 5 spikes were detected per minute. A burst was considered as a group of at least 5 spikes, each separated by an inter-spike interval (ISI) of no more than 100 ms. Network bursts were identified as a minimum of 10 spikes with a maximum ISI of 100 milliseconds covered by at least 25% of electrodes in each well. No non-active well was excluded in the analysis. After the last reading at week 8, selected wells were treated with three synaptic antagonists: GABA_A_ receptor antagonist picrotoxin (PTX; Sigma) at 100 μM, AMPA receptor antagonist 6-cyano-7-nitroquinoxaline-2,3-dion (CNQX; Sigma) at 60 μM, and sodium ion channel antagonist tetrodotoxin (TTX; Alomone labs) at 1 μM. The plates were recorded consecutively, 5-10 minutes after addition of the antagonists. A 60-minute recovery period was allowed in the incubator at 37°C between each antagonist treatment and plate recording.

## References

Anagnostou, E., Zwaigenbaum, L., Szatmari, P., Fombonne, E., Fernandez, B.A., Woodbury-Smith, M., Brian, J., Bryson, S., Smith, I.M., Drmic, I., et al. (2014). Autism spectrum disorder: advances in evidence-based practice. CMAJ 186, 509–519.

Bakkaloglu, B., O’Roak, B.J., Louvi, A., Gupta, A.R., Abelson, J.F., Morgan, T.M., Chawarska, K., Klin, A., Ercan-Sencicek, A.G., Stillman, A.A., et al. (2008). Molecular cytogenetic analysis and resequencing of contactin associated protein-like 2 in autism spectrum disorders. Am J Hum Genet 82, 165–173.

Bensaid, M., Melko, M., Bechara, E.G., Davidovic, L., Berretta, A., Catania, M.V., Gecz, J., Lalli, E., and Bardoni, B. (2009). FRAXE-associated mental retardation protein (FMR2) is an RNA-binding protein with high affinity for G-quartet RNA forming structure. Nucleic Acids Res 37, 1269–1279.

Bernier, R., Golzio, C., Xiong, B., Stessman, H.A., Coe, B.P., Penn, O., Witherspoon, K., Gerdts, J., Baker, C., Vulto-van Silfhout, A.T., et al. (2014). Disruptive CHD8 mutations define a subtype of autism early in development. Cell 158, 263–276.

Betancur, C. (2011). Etiological heterogeneity in autism spectrum disorders: more than 100 genetic and genomic disorders and still counting. Brain Res 1380, 42–77.

Bourgeron, T. (2015). From the genetic architecture to synaptic plasticity in autism spectrum disorder. Nat Rev Neurosci 16, 551–563.

Brett, M., McPherson, J., Zang, Z.J., Lai, A., Tan, E.S., Ng, I., Ong, L.C., Cham, B., Tan, P., Rozen, S., et al. (2014). Massively parallel sequencing of patients with intellectual disability, congenital anomalies and/or autism spectrum disorders with a targeted gene panel. PLoS One 9, e93409.

Carter, M.T., and Scherer, S.W. (2013). Autism spectrum disorder in the genetics clinic: a review. Clin Genet 83, 399–407.

Colvert, E., Tick, B., McEwen, F., Stewart, C., Curran, S.R., Woodhouse, E., Gillan, N., Hallett, V., Lietz, S., Garnett, T., et al. (2015). Heritability of Autism Spectrum Disorder in a UK Population-Based Twin Sample. JAMA Psychiatry 72, 415–423.

de la Torre-Ubieta, L., Won, H., Stein, J.L., and Geschwind, D.H. (2016). Advancing the understanding of autism disease mechanisms through genetics. Nat Med 22, 345–361.

De Rubeis, S., He, X., Goldberg, A.P., Poultney, C.S., Samocha, K., Cicek, A.E., Kou, Y., Liu, L., Fromer, M., Walker, S., et al. (2014). Synaptic, transcriptional and chromatin genes disrupted in autism. Nature 515, 209–215.

Djuric, U., Rodrigues, D.C., Batruch, I., Ellis, J., Shannon, P., and Diamandis, P. (2017). Spatiotemporal Proteomic Profiling of Human Cerebral Development. Mol Cell Proteomics 16, 1548–1562.

DSM-V (2013). Diagnostic and Statistical Manual of Mental Disorders, Fifth Edition. American Psychiatric Association.

Durak, O., Gao, F., Kaeser-Woo, Y.J., Rueda, R., Martorell, A.J., Nott, A., Liu, C.Y., Watson, L.A., and Tsai, L.H. (2016). Chd8 mediates cortical neurogenesis via transcriptional regulation of cell cycle and Wnt signaling. Nat Neurosci 19, 1477–1488.

Geschwind, D.H., and State, M.W. (2015). Gene hunting in autism spectrum disorder: on the path to precision medicine. Lancet Neurol 14, 1109–1120.

Gilman, S.R., Iossifov, I., Levy, D., Ronemus, M., Wigler, M., and Vitkup, D. (2011). Rare de novo variants associated with autism implicate a large functional network of genes involved in formation and function of synapses. Neuron 70, 898–907.

Gronborg, T.K., Schendel, D.E., and Parner, E.T. (2013). Recurrence of autism spectrum disorders in full- and half-siblings and trends over time: a population-based cohort study. JAMA Pediatr 167, 947–953.

Habela, C.W., Song, H., and Ming, G.L. (2016). Modeling synaptogenesis in schizophrenia and autism using human iPSC derived neurons. Mol Cell Neurosci 73, 52–62.

Hardelin, J.P., Levilliers, J., Blanchard, S., Carel, J.C., Leutenegger, M., Pinard-Bertelletto, J.P., Bouloux, P., and Petit, C. (1993). Heterogeneity in the mutations responsible for X chromosome-linked Kallmann syndrome. Hum Mol Genet 2, 373–377.

Hirano, S., and Takeichi, M. (2012). Cadherins in brain morphogenesis and wiring. Physiol Rev 92, 597–634.

Hotta, A., Cheung, A.Y., Farra, N., Vijayaragavan, K., Seguin, C.A., Draper, J.S., Pasceri, P., Maksakova, I.A., Mager, D.L., Rossant, J., et al. (2009). Isolation of human iPS cells using EOS lentiviral vectors to select for pluripotency. Nat Methods 6, 370–376.

Hsu, P.D., Scott, D.A., Weinstein, J.A., Ran, F.A., Konermann, S., Agarwala, V., Li, Y., Fine, E.J., Wu, X., Shalem, O., et al. (2013). DNA targeting specificity of RNA-guided Cas9 nucleases. Nat Biotechnol 31, 827–832.

Hussman, J.P. (2001). Suppressed GABAergic inhibition as a common factor in suspected etiologies of autism. J Autism Dev Disord 31, 247–248.

Jiang, Y.H., Yuen, R.K., Jin, X., Wang, M., Chen, N., Wu, X., Ju, J., Mei, J., Shi, Y., He, M., et al. (2013). Detection of clinically relevant genetic variants in autism spectrum disorder by whole-genome sequencing. Am J Hum Genet 93, 249–263.

Jung, E.M., Moffat, J.J., Liu, J., Dravid, S.M., Gurumurthy, C.B., and Kim, W.Y. (2017). Arid1b haploinsufficiency disrupts cortical interneuron development and mouse behavior. Nat Neurosci 20, 1694–1707.

Kamiya, K., Kaneda, M., Sugawara, T., Mazaki, E., Okamura, N., Montal, M., Makita, N., Tanaka, M., Fukushima, K., Fujiwara, T., et al. (2004). A nonsense mutation of the sodium channel gene SCN2A in a patient with intractable epilepsy and mental decline. J Neurosci 24, 2690–2698.

Kim, H.J., and Magrane, J. (2011). Isolation and culture of neurons and astrocytes from the mouse brain cortex. Methods Mol Biol 793, 63–75.

Kumar, R.A., KaraMohamed, S., Sudi, J., Conrad, D.F., Brune, C., Badner, J.A., Gilliam, T.C., Nowak, N.J., Cook, E.H., Jr., Dobyns, W.B., et al. (2008). Recurrent 16p11.2 microdeletions in autism. Hum Mol Genet 17, 628–638.

Kuratomi, G., Iwamoto, K., Bundo, M., Kusumi, I., Kato, N., Iwata, N., Ozaki, N., and Kato, T. (2008). Aberrant DNA methylation associated with bipolar disorder identified from discordant monozygotic twins. Mol Psychiatry 13, 429–441.

Lai, M.C., Lombardo, M.V., and Baron-Cohen, S. (2014). Autism. Lancet 383, 896–910.

Law, M.J., Lower, K.M., Voon, H.P., Hughes, J.R., Garrick, D., Viprakasit, V., Mitson, M., De Gobbi, M., Marra, M., Morris, A., et al. (2010). ATR-X syndrome protein targets tandem repeats and influences allele-specific expression in a size-dependent manner. Cell 143, 367–378.

Lee, S.G., Joo, Y., Kim, B., Chung, S., Kim, H.L., Lee, I., Choi, B., Kim, C., and Song, K. (2005). Association of Ala72Ser polymorphism with COMT enzyme activity and the risk of schizophrenia in Koreans. Hum Genet 116, 319–328.

Lionel, A.C., Tammimies, K., Vaags, A.K., Rosenfeld, J.A., Ahn, J.W., Merico, D., Noor, A., Runke, C.K., Pillalamarri, V.K., Carter, M.T., et al. (2014). Disruption of the ASTN2/TRIM32 locus at 9q33.1 is a risk factor in males for autism spectrum disorders, ADHD and other neurodevelopmental phenotypes. Hum Mol Genet 23, 2752–2768.

Liu, H., Chang, L.H., Sun, Y., Lu, X., and Stubbs, L. (2014). Deep vertebrate roots for mammalian zinc finger transcription factor subfamilies. Genome Biol Evol 6, 510–525.

Luo, H., Zheng, R., Zhao, Y., Wu, J., Li, J., Jiang, F., Chen, D.N., Zhou, X.T., and Li, J.D. (2017). A dominant negative FGFR1 mutation identified in a Kallmann syndrome patient. Gene 621, 1–4.

Marshall, C.R., Noor, A., Vincent, J.B., Lionel, A.C., Feuk, L., Skaug, J., Shago, M., Moessner, R., Pinto, D., Ren, Y., et al. (2008). Structural variation of chromosomes in autism spectrum disorder. Am J Hum Genet 82, 477–488.

McKenna, A., Hanna, M., Banks, E., Sivachenko, A., Cibulskis, K., Kernytsky, A., Garimella, K., Altshuler, D., Gabriel, S., Daly, M., et al. (2010). The Genome Analysis Toolkit: a MapReduce framework for analyzing next-generation DNA sequencing data. Genome Res 20, 1297–1303.

Miyaoka, Y., Chan, A.H., Judge, L.M., Yoo, J., Huang, M., Nguyen, T.D., Lizarraga, P.P., So, P.L., and Conklin, B.R. (2014). Isolation of single-base genome-edited human iPS cells without antibiotic selection. Nat Methods 11, 291–293.

Moretto, E., Murru, L., Martano, G., Sassone, J., and Passafaro, M. (2017). Glutamatergic synapses in neurodevelopmental disorders. Prog Neuropsychopharmacol Biol Psychiatry.

Nishiyama, M., Nakayama, K., Tsunematsu, R., Tsukiyama, T., Kikuchi, A., and Nakayama, K.I. (2004). Early embryonic death in mice lacking the beta-catenin-binding protein Duplin. Mol Cell Biol 24, 8386–8394.

Ozonoff, S., Young, G.S., Carter, A., Messinger, D., Yirmiya, N., Zwaigenbaum, L., Bryson, S., Carver, L.J., Constantino, J.N., Dobkins, K., et al. (2011). Recurrence risk for autism spectrum disorders: a Baby Siblings Research Consortium study. Pediatrics 128, e488–495.

Pak, C., Danko, T., Zhang, Y., Aoto, J., Anderson, G., Maxeiner, S., Yi, F., Wernig, M., and Sudhof, T.C. (2015). Human Neuropsychiatrie Disease Modeling using Conditional Deletion Reveals Synaptic Transmission Defects Caused by Heterozygous Mutations in NRXN1. Cell Stem Cell 17, 316–328.

Pinto, D., Delaby, E., Merico, D., Barbosa, M., Merikangas, A., Klei, L., Thiruvahindrapuram, B., Xu, X., Ziman, R., Wang, Z., et al. (2014). Convergence of genes and cellular pathways dysregulated in autism spectrum disorders. Am J Hum Genet 94, 677–694.

Pinto, D., Pagnamenta, A.T., Klei, L., Anney, R., Merico, D., Regan, R., Conroy, J., Magalhaes, T.R., Correia, C., Abrahams, B.S., et al. (2010). Functional impact of global rare copy number variation in autism spectrum disorders. Nature 466, 368–372.

Planells-Cases, R., Caprini, M., Zhang, J., Rockenstein, E.M., Rivera, R.R., Murre, C., Masliah, E., and Montal, M. (2000). Neuronal death and perinatal lethality in voltagegated sodium channel alpha(II)-deficient mice. Biophys J 78, 2878–2891.

Platt, R.J., Zhou, Y., Slaymaker, I.M., Shetty, A.S., Weisbach, N.R., Kim, J.A., Sharma, J., Desai, M., Sood, S., Kempton, H.R., et al. (2017). Chd8 Mutation Leads to Autistic-like Behaviors and Impaired Striatal Circuits. Cell Rep 19, 335–350.

Ran, F.A., Hsu, P.D., Lin, C.Y., Gootenberg, J.S., Konermann, S., Trevino, A.E., Scott, D.A., Inoue, A., Matoba, S., Zhang, Y., et al. (2013). Double nicking by RNA-guided CRISPR Cas9 for enhanced genome editing specificity. Cell 154, 1380–1389.

Risch, N., Hoffmann, T.J., Anderson, M., Croen, L.A., Grether, J.K., and Windham, G.C. (2014). Familial recurrence of autism spectrum disorder: evaluating genetic and environmental contributions. Am J Psychiatry 171, 1206–1213.

Rubenstein, J.L., and Merzenich, M.M. (2003). Model of autism: increased ratio of excitation/inhibition in key neural systems. Genes Brain Behav 2, 255–267.

Sahin, M., and Sur, M. (2015). Genes, circuits, and precision therapies for autism and related neurodevelopmental disorders. Science 350.

Takahashi, K., Tanabe, K., Ohnuki, M., Narita, M., Ichisaka, T., Tomoda, K., and Yamanaka, S. (2007). Induction of pluripotent stem cells from adult human fibroblasts by defined factors. Cell 131, 861–872.

Tammimies, K., Marshall, C.R., Walker, S., Kaur, G., Thiruvahindrapuram, B., Lionel, A.C., Yuen, R.K., Uddin, M., Roberts, W., Weksberg, R., et al. (2015). Molecular Diagnostic Yield of Chromosomal Microarray Analysis and Whole-Exome Sequencing in Children With Autism Spectrum Disorder. JAMA 314, 895–903.

Varghese, M., Keshav, N., Jacot-Descombes, S., Warda, T., Wicinski, B., Dickstein, D.L., Harony-Nicolas, H., De Rubeis, S., Drapeau, E., Buxbaum, J.D., et al. (2017). Autism spectrum disorder: neuropathology and animal models. Acta Neuropathol 134, 537–566.

Weiss, L.A., Shen, Y., Korn, J.M., Arking, D.E., Miller, D.T., Fossdal, R., Saemundsen, E., Stefansson, H., Ferreira, M.A., Green, T., et al. (2008). Association between microdeletion and microduplication at 16p11.2 and autism. N Engl J Med 358, 667–675.

Wintle, R.F., Lionel, A.C., Hu, P., Ginsberg, S.D., Pinto, D., Thiruvahindrapduram, B., Wei, J., Marshall, C.R., Pickett, J., Cook, E.H., et al. (2011). A genotype resource for postmortem brain samples from the Autism Tissue Program. Autism Res 4, 89–97.

Yang, L., Mali, P., Kim-Kiselak, C., and Church, G. (2014). CRISPR-Cas-mediated targeted genome editing in human cells. Methods Mol Biol 1114, 245–267.

Yang, N., Chanda, S., Marro, S., Ng, Y.H., Janas, J.A., Haag, D., Ang, C.E., Tang, Y., Flores, Q., Mall, M., et al. (2017). Generation of pure GABAergic neurons by transcription factor programming. Nat Methods 14, 621–628.

Yi, F., Danko, T., Botelho, S.C., Patzke, C., Pak, C., Wernig, M., and Sudhof, T.C. (2016). Autism-associated SHANK3 haploinsufficiency causes Ih channelopathy in human neurons. Science 352, aaf2669.

Yu, J., Vodyanik, M.A., Smuga-Otto, K., Antosiewicz-Bourget, J., Frane, J.L., Tian, S., Nie, J., Jonsdottir, G.A., Ruotti, V., Stewart, R., et al. (2007). Induced pluripotent stem cell lines derived from human somatic cells. Science 318, 1917–1920.

Yuen, R.K., Merico, D., Bookman, M., J, L.H., Thiruvahindrapuram, B., Patel, R.V., Whitney, J., Deflaux, N., Bingham, J., Wang, Z., et al. (2017). Whole genome sequencing resource identifies 18 new candidate genes for autism spectrum disorder. Nat Neurosci 20, 602–611.

Yuen, R.K., Merico, D., Cao, H., Pellecchia, G., Alipanahi, B., Thiruvahindrapuram, B., Tong, X., Sun, Y., Cao, D., Zhang, T., et al. (2016). Genome-wide characteristics of de novo mutations in autism. NPJ Genom Med 1, 160271–1602710.

Yuen, R.K., Thiruvahindrapuram, B., Merico, D., Walker, S., Tammimies, K., Hoang, N., Chrysler, C., Nalpathamkalam, T., Pellecchia, G., Liu, Y., et al. (2015). Whole-genome sequencing of quartet families with autism spectrum disorder. Nat Med 21, 185–191.

Zhang, Y., Pak, C., Han, Y., Ahlenius, H., Zhang, Z., Chanda, S., Marro, S., Patzke, C., Acuna, C., Covy, J., et al. (2013). Rapid single-step induction of functional neurons from human pluripotent stem cells. Neuron 78, 785–798.

